# Exploring per-base quality scores as a surrogate marker of cell-free DNA fragmentome

**DOI:** 10.64898/2026.03.08.710357

**Authors:** Hadas Volkov, Maria Raitses-Gurevich, Meitar Grad, Rani Shlayem, Dev Leibowitz, Tami Rubinek, Talia Golan, Noam Shomron

**Author notes:** Correspondence to: Hadas Volkov, Noam Shomron.

## Abstract

Per-base quality scores are widely treated as technical metadata in next-generation sequencing. Here, we show that in rigorously controlled whole-genome sequencing of cell-free DNA, quality profiles may encode fragmentomic signals that enable classification of cancer samples against matched controls. Analyzing four independent batches (23 cancer samples: pancreatic and breast; 22 matched controls) sequenced in a within-lane regime and further normalized per flow-cell tile to reduce technical confounders, we demonstrate through unsupervised analysis that boundary-enriched dynamics captured in these quality scores consistently separate cancer from control samples. A leave-one-batch-out classifier trained on quality-derived scores achieved a pooled area under the curve of 0.81. Furthermore, we show that the quality-derived metric correlates with short-fragment enrichment and tumor-associated 5’-end motifs, performing comparably to established, motif-based orthogonal methods. These results provide initial evidence that quality scores could serve as a low-cost, alignment-free biomarker for cfDNA-based cancer detection.

**Key Points:**

- PBQS in rigorously controlled cfDNA whole-genome sequencing contain biologically informative fragmentomic signal rather than only technical noise
- Boundary-enriched quality dynamics distinguish cancer samples from matched controls across independent sequencing batches
- A leave-one-batch-out classifier based on PBQS-derived features achieved a pooled AUC of 0.81 across 23 cancer and 22 control samples
- The PBQS-derived score correlates with short-fragment enrichment and tumor-associated 5′ end motifs, supporting its value as a lightweight orthogonal biomarker for cfDNA cancer.

**Biographical Note:** Prof. Noam Shomron heads the Functional Genomics Laboratory at Tel Aviv University’s Medical School, where his group studies genomics and bioinformatics with a focus on sequencing technologies and translational medicine.

## Introduction

Next-generation sequencing (NGS) technologies have revolutionized biological research, enabling unprecedented insights into the genome, transcriptome, and epigenome. A fundamental output of NGS is the sequence of nucleotide bases for each sequenced fragment, accompanied by a crucial metric known as the per-base quality scores (PBQS). These scores, typically represented as Phred scores^1,2^, provide a probabilistic assessment of the accuracy of each base call. Specifically, a Phred score (Q) is logarithmically related to the probability of an incorrect base call (*P*), according to the formula

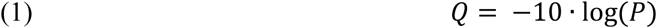

Consequently, higher Phred scores indicate a lower probability of error and greater confidence in the base call.

The importance of PBQS in downstream NGS data analysis cannot be overstated; they serve as a cornerstone for quality control procedures, informing decisions on read trimming to remove low-quality bases, thereby improving the accuracy of subsequent analyses^3^. Furthermore, PBQS are integral to variant calling algorithms^4,5^, where they are used to weight the evidence for each potential variant, with lower quality bases contributing less to the overall confidence in a variant call. Beyond individual reads and variants, aggregated base quality metrics are also used to assess the overall quality of a sequencing run^6,7^.

Traditionally, variations in PBQS are primarily attributed to technical factors inherent in the NGS process. These factors can be broadly categorized and include platform-specific characteristics of the sequencing instrument^8^. For instance, on Illumina platforms, which utilize sequencing-by-synthesis (SBS) chemistry, error modes such as phasing and pre-phasing can influence base calling accuracy and consequently, quality scores^9^. The quality and consistency of reagents, as well as the precise calibration of the sequencing instrument’s optics and fluidics, also play a significant role. Furthermore, the library preparation process itself can introduce biases and artifacts that impact base qualities. PCR amplification, a common step in many library preparation protocols, can introduce errors^10^ and biases that manifest in the sequencing data. Similarly, the efficiency of adapter ligation and the chosen library size selection protocols can influence the signal-to-noise ratio and ultimately, base quality scores. Sequencing run parameters, such as cluster density on the flow cell and read length, can also affect the fidelity of base calls. Finally, the algorithms used for base calling during primary data processing are crucial determinants of the final quality scores assigned to each base.

The prevailing understanding within the field is that PBQS are predominantly a technical metric, reflecting the performance and inherent limitations of the sequencing technology and associated protocols^8,10^. Variations in these scores are typically interpreted as indicators of technical artifacts or inconsistencies in the experimental process. Nevertheless, biological processes such as DNA secondary structures, Guanine-Cytosine (GC) content bias, and fragment length may significantly and systematically impact PBQS in Illumina sequencing by introducing sequencing errors, signal decay, and polymerization inefficiencies, with GC bias^11,12^, non-B DNA structures^13,14^ and fragment length^15^ being the most consistently identified contributors described and investigated^16^. Although no evidence connects cancer-specific genomic changes directly to PBQS profile changes, available literature highlights the compounding effects of systematic errors, mutational overrepresentation, copy number variations (CNVs), and sequencing biases on Illumina workflows in cancer samples^17,18^.

While PBQS have been predominantly attributed to technical factors inherent in the NGS workflow, the characteristics of cell-free DNA (cfDNA) fragments, collectively referred to as fragmentomics, along with cancer-associated genetic and epigenetic modifications, may systematically impact PBQS profiles. cfDNA originates from apoptotic and necrotic like cell processes^19^, resulting in DNA fragments that exhibit distinct size distributions and nucleotide patterns compared to genomic DNA extracted from tissues or cells. These unique fragmentomic features have been exploited for non-invasive cancer diagnostics and monitoring, as they can reflect the underlying tumor biology and the presence of malignancy^20–22^.

Cancer-related modifications, such as somatic mutations, CNVs, and aberrant methylation patterns, alter the genomic landscape and are hallmarks of cancer. Nevertheless, we speculated that the physical properties of cfDNA fragments, driven by nuclease activity, may play a more direct role in influencing the efficiency of the sequencing reaction when it comes to cfDNA. Characteristic shifts in fragmentation patterns between cfDNA shed from non-cancer cells and the tumor-derived cfDNA (ctDNA) fraction have been investigated and utilized extensively for cancer monitoring^20,23^. These features include shorter ctDNA fragments, specific 5’ and 3’ end-motifs and potentially jagged single-stranded overhangs among others. In the context of SBS, these specific end-motif sequences and length differences may challenge the sequencing chemistry. For example, over-representation of specific nucleotide motif at fragment end might be more prone to phasing or pre-phasing errors or affect polymerase kinetics during specific sequencing cycles, thereby systematically altering the assigned quality scores.

Given these considerations, we hypothesize that the fragmentomic features inherent to cfDNA from cancer patients can systematically influence PBQS profiles. We propose that PBQS is not merely a record of technical noise but rather acts as a latent surrogate marker for underlying fragmentomic properties. We anticipate that this influence manifests as distinct quality score patterns, particularly at the read boundaries where fragmentomic signatures are most pronounced. Disentangling this biological signal from technical artifacts could allow raw quality scores to serve as a computationally efficient, orthogonal biomarker for cancer detection.

## Results

### Unsupervised PBQS analysis suggests cancer stratification in cfDNA

To test the hypothesis that PBQS profiles harbor latent fragmentomic information, we assembled a multi-center dataset designed intentionally at minimizing technical confounders. The study comprised four independent sequencing batches, including patients with pancreatic ductal adenocarcinoma (PDAC), breast cancer, and their respective technical matched controls (Fig. 1a). Crucially, to mitigate technical artifacts at the physical sequencing level, cancer and control samples within each batch were handled simultaneously and sequenced on a single flow cell lane. The PDAC cohort consisted of three batches derived from two different medical centers while sharing a uniform library preparation and sequencing regime (paired end, 151 bp, Table S1).

**Fig. 1:**
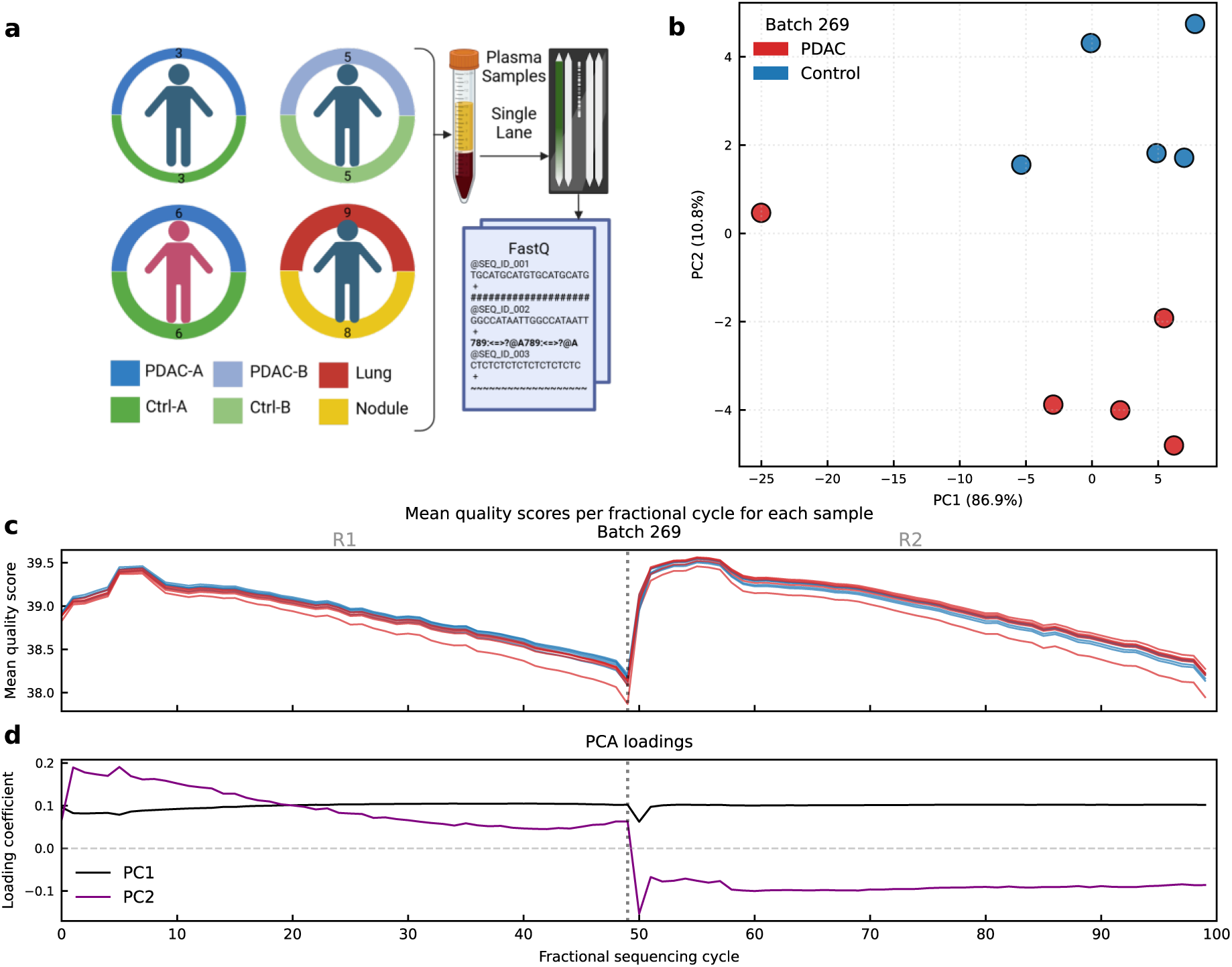
Unsupervised analysis of PBQS suggests cancer stratification. **a** Schematic overview of the study cohort and experimental design. The dataset comprises four independent sequencing batches: three involving PDAC patients and one involving breast cancer patients, alongside matched controls (healthy donors or benign findings). Batch 269 (PDAC 5/5), Batch 102 (PDAC 3/3), Batch 42 (PDAC 6/6), and Batch 149 (breast cancer/benign 9/8), totaling 23 cancer and 22 control samples. To minimize technical confounders, cancer and control samples within each batch were processed simultaneously and sequenced on a single flow cell lane. **b** PCA projection of a representative PDAC batch (Batch 269). The scatter plot reveals a distinct separation between cancer samples (red) and controls (blue) along the second principal component (PC2), despite PC1 capturing the majority of the variance. **c** MFPQP for Batch 269. The traces display the aggregated per-base quality scores mapped to a standardized fractional vector (*K* = 100, concatenating Read 1 and Read 2). Cancer samples exhibit a subtle but systematic elevation in quality scores compared to controls. **d** PCA loading vectors for the first two principal components. The PC1 loading vector (black) remains relatively flat across the sequencing cycle, indicating it tracks global mean quality shifts likely attributable to technical variation. In contrast, the PC2 loading vector (purple) exhibits distinct dynamic fluctuations concentrated at the read boundaries (5’ and 3’ ends), aligning with the hypothesis that PC2 captures latent fragmentomic features.

To validate the generalizability of the approach, we included a fourth independent batch of breast cancer patients processed at a separate facility with distinct sample preprocessing chemistries and a dissimilar sequencing regime (paired end, 143 bp). Notably, cancer patients in the cohort consisted of stage I-II with small primary tumors (pT1–pT2) and minimal nodal involvement (including micrometastases, pN1(mi)). Such early-stage localized cancers are widely characterized as ‘low-shedding’ phenotypes, typically yielding low tumor fraction^24^. Furthermore, the control group for the cohort consisted of women with benign findings (e.g., fibroadenomas) (Table S2). Across the four sequencing batches, sample sizes (cancer/control) were: Batch 269 (PDAC 5/5), Batch 102 (PDAC 3/3), Batch 42 (PDAC 6/6), and Batch 149 (breast cancer/benign 9/8), totaling 23 cancer and 22 control samples (*N* = 45).

To further reduce the possibility of technical noise propagating into the quality profiles, we applied a preprocessing pipeline that enforced equal read amounts per flow-cell tile. This step was critical to avoid quality shifts driven by variations in cluster density or coverage bias^25,26^. To accommodate variable read lengths across batches and samples, we computed the Mean Fractional Position Quality Profile (MFPQP). This method maps variable-length reads to a standardized vector (*K* = 100, representing concatenated R1 and R2), allowing for the calculation of a comparable mean per-base quality vector for every sample (Methods).

We initially visualized the MFPQP of a representative PDAC batch (Batch 269) to assess raw signal differences. Visual inspection reveals a subtle systematic bias along the profile, where cancer-derived samples exhibit slightly higher quality scores compared to controls (Fig. 1c). To decompose this variation, we applied an unsupervised Principal Component Analysis (PCA). The resulting projection reveals a clear separation between the groups (Fig. 1b). The first Principal Component (PC1) accounts for the majority of the variance (86.9%), while PC2 accounts for 10.8%. However, PC2 defines the axis of separation between cancer and control samples. Similar patterns emerge for each batch in the dataset (Fig. S1-4). Inspection of the component loadings (Fig. 1d) provides a biological intuition for the mathematical separation. The PC1 loadings remain relatively flat across sequencing cycles suggesting that PC1 tracks the overall magnitude due to collinearity of the MFPQP values across samples, thus serves as a summary of the shared variance^27^. In contrast, PC2 loadings exhibit distinct dynamics concentrated at the beginning of the reads. This variance profile, at the 5’ and 3’ ends, aligns with regions where fragmentomic features (i.e., end-motifs and fragment lengths) are pronounced, thus giving rise to the assumption that PC2 captures a biological signal and PC1 acts as a proxy for total intensity.

### PBQS profiles harbor batch-independent cancer signature

To further generalize the PBQS stratification observed in a single batch analysis we expanded the analysis to the complete dataset. We aggregated the quality profiles from all four batches into a unified analysis while applying Z-score transformation to each batch independently to overcome batch specific quality magnitudes (PCA projection Figure 2a). Consistent with our initial single-batch observations, PC1 captures the largest share of variance (82.7%), yet it shows negligible association with the biological label (sample-size weighted Cohen’s *d* = 0.02, Table S3). In contrast, PC2 consistently separated the cancer samples from matched technical controls within each individual batch (Fig. 2b), and exhibiting a strong effect size (Cohen’s *d* = 1.19, Table S3). To formally quantify this independence, a fitted Ordinary Least Squares (OLS) regression model (*PC*2 ∼ *Label* + *Batch*) indicates that the cancer label remains a significant predictor of the PC2 score (*P* = 0.009) while adjusting for batch identity (Table S4).

**Fig. 2:**
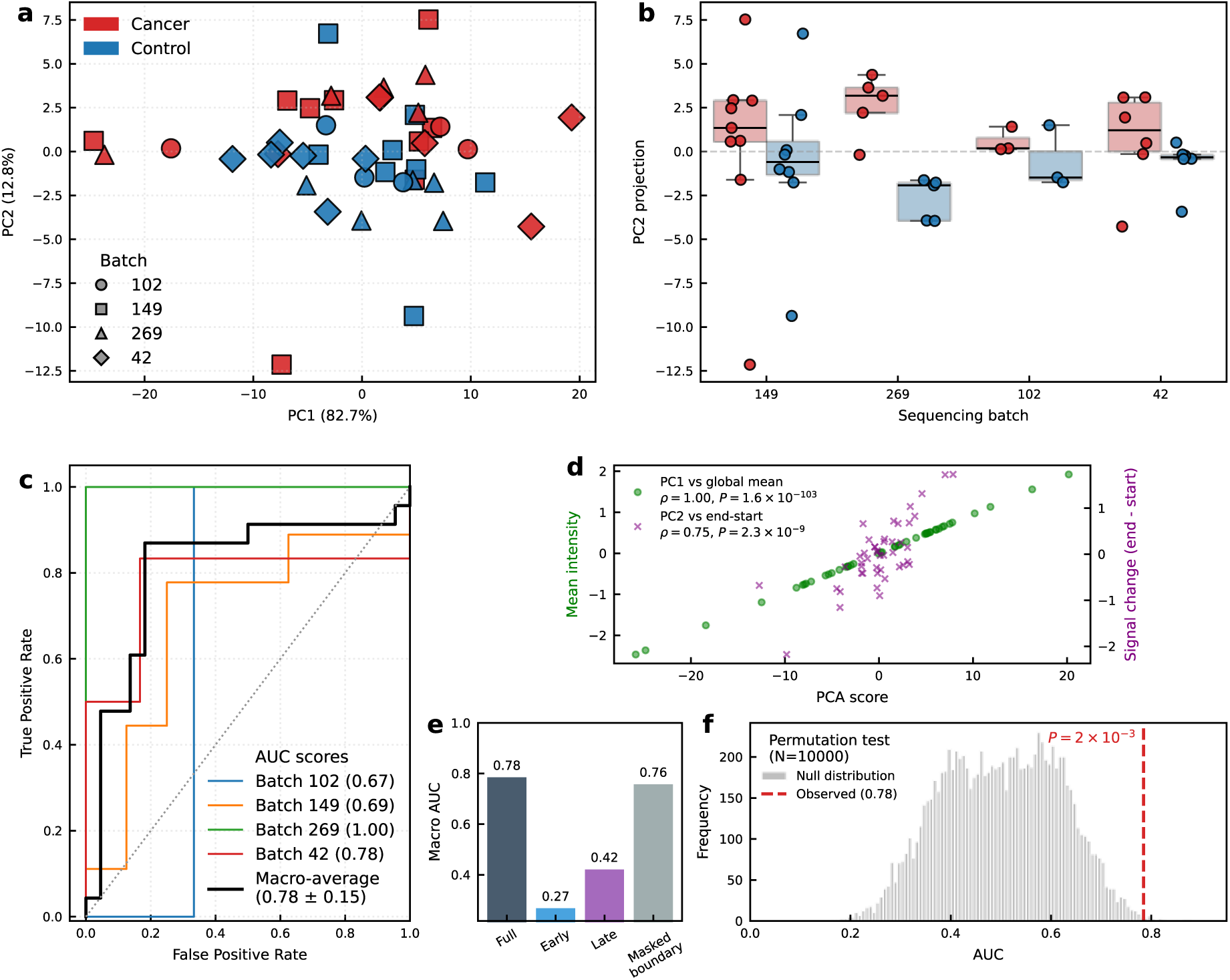
PBQS profiles harbor batch-independent cancer signature. **a** PCA projection of the aggregated per batch Z-scored dataset shaped by sequencing batch (red: cancer, blue: control). While PC1 captures the majority of variance (82.7%), PC2 (12.8%) defines the axis of separation between cancer and control samples. **b** Distribution of PC2 scores stratified by sequencing batch. In all four independent batches, cancer samples exhibit consistently higher PC2 scores compared to their matched technical controls, confirming the directionality of the signal is not batch dependent. **c** ROC curves from LOBO cross-validation. Colored lines indicate performance when testing on each specific held-out batch, while the black line represents the macro-average performance (AUC = 0.78 ± 0.15) **d** Correlation analysis linking latent PCA components to physical quality attributes. PC1 (green circles) shows perfect correlation with the global mean quality score (Pearson’s ρ = 1.00), indicating it tracks overall sequencing magnitude. PC2 (purple crosses) correlates significantly with the “end-start” quality delta (Pearson’s ρ = 0.75), suggesting it captures the dynamic slope of quality evolution across the read. **e** Feature ablation study comparing LOBO performance across different segments of the quality profile. The “Masked boundary” model, which retains only the 5’ and 3’ termini, preserves classification power (AUC = 0.76), whereas models restricted to only the early or late segments perform at or below random chance. **f** Statistical significance assessment via stratified permutation testing (*n* = 10,000 iterations). The histogram displays the null distribution of AUC scores generated by randomly shuffling labels within batches. The observed AUC (red dashed line, 0.78) is statistically significant (*P* = 0.002)

To evaluate transferability among batches, we implemented a Leave-One-Batch-Out (LOBO) cross-validation scheme. In this approach, a model is trained on three batches and tested on the unseen fourth batch, thus facilitating classification performance assessment. The PBQS PC2-based classifier achieved a global pooled Area Under the Curve (AUC) of 0.81 and a macro-average AUC of 0.78 ± 0.15 across the four folds (Fig. 2c, Table S5). Notably, the model successfully classified samples from the breast cancer batch (Batch 149, AUC = 0.69) despite being trained exclusively on PDAC samples. This is significant given that this batch comprised of low ctDNA shedding samples, which typically poses a detection challenge. Lastly, we achieved full separation in one of the PDAC batches (Batch 269, AUC = 1.00). This robust performance likely reflects the combination of higher sequencing depth and the inclusion of more advanced disease stages, which likely enhance the fragmentomic-derived quality signature (Table S1, Table S8).

In an attempt to investigate mechanistic relation to fragmentomic features we sought to decouple the physical attributes of the quality profiles from the PCA latent representation. Correlation analysis revealed that PC1 is fully correlated with the global mean quality score (Pearson’s ρ = 1.00, *P* < 10^−100^), strengthening the assumption that it acts as a proxy for overall magnitude (Fig. 2d, Fig. S5). Conversely, PC2 strongly correlates with the “end-start” quality delta (ρ = 0.75, *P* < 10^−8^), representing the slope or dynamic change in quality across the read. This supports the notion that the cancer signal is encoded in the dynamic evolution of quality scores along the fragment rather than the absolute quality value.

To further localize this signal, we performed an ablation study (Fig. 2e). By masking the inner regions of the PBQS vector and keeping the 5’ and 3’ ends of the reads (“Masked Boundary” features), the model retained the vast majority of the classification power (Macro AUC = 0.76, Table S6). In contrast, models trained solely on the early or late segments of the reads dropped below random performance (Table S6), suggesting that the cancer-specific signal is physically concentrated at the fragment termini. Although early-cycle effects should be shared across labels due to within-lane sequencing and tile-level normalization, we additionally tested whether the signal depended on the first few bases. Removing the first 3, 5, or 10 bases from both reads did not reduce LOBO performance and fixed 10 bp absolute-cycle binning showed that the most predictive bins were distributed beyond the first 10 bp of either read (Table S7).

Additionally, to ensure that these results were not a product of random chance or class imbalance, we performed a stratified permutation test (Fig. 2f). By randomly shuffling labels within batches over 10,000 iterations, we established a null distribution of LOBO performance. The observed macro-average AUC of 0.78 fell significantly outside the null distribution (*P* = 0.002), establishing statistical significance for the PC2-based classifier.

Lastly, because MFPQP maps reads of different lengths onto a shared fractional coordinate system, we tested whether the observed separation could be attributed to the fractional-position representation itself. We repeated the analysis using absolute sequencing-cycle coordinates in the three PDAC batches, all of which were generated under the same 2×151 bp sequencing regime. In a representative batch, the absolute-cycle quality profiles and PCA loadings recapitulated the qualitative structure observed with MFPQP, including separation along the second principal component (Fig. S6). Across all three PDAC batches, the absolute-cycle representation retained discrimination in the LOBO framework, with a macro-average AUC of 0.79 ± 0.18 (Fig. S7). These results indicate that the PBQS signal is not solely an artifact of fractional-position mapping. Rather, MFPQP serves primarily as a harmonization strategy for integrating batches with different read lengths.

### PBQS derived score correlate with fragmentomic features

Having suggested that technically controlled PBQS derived scores may stratify cancer samples from controls, we sought to offer plausible biological mechanism allowing this. We assumed that distinct size distributions and end-motifs characteristics of ctDNA are leading factors. Consistent with established literature; cancer samples in our dataset exhibited the common enrichment of shorter fragments surrounding the typical mononucleosomal peak (∼167 bp) and a depletion of longer fragments compared to healthy controls (Fig. 3a). To assess if these physical length differences might influence sequencing quality, we binned MFPQPs by their respective fragment lengths and computed per cycle means. This analysis demonstrates a visual relationship where overall per cycle quality magnitudes decrease as fragment length increases (representative Batch 269, Fig. 3c), suggesting an association as previously described by Tan *et al.,* linking long fragments to reduced quality in Illumina SBS^15^.

**Fig. 3:**
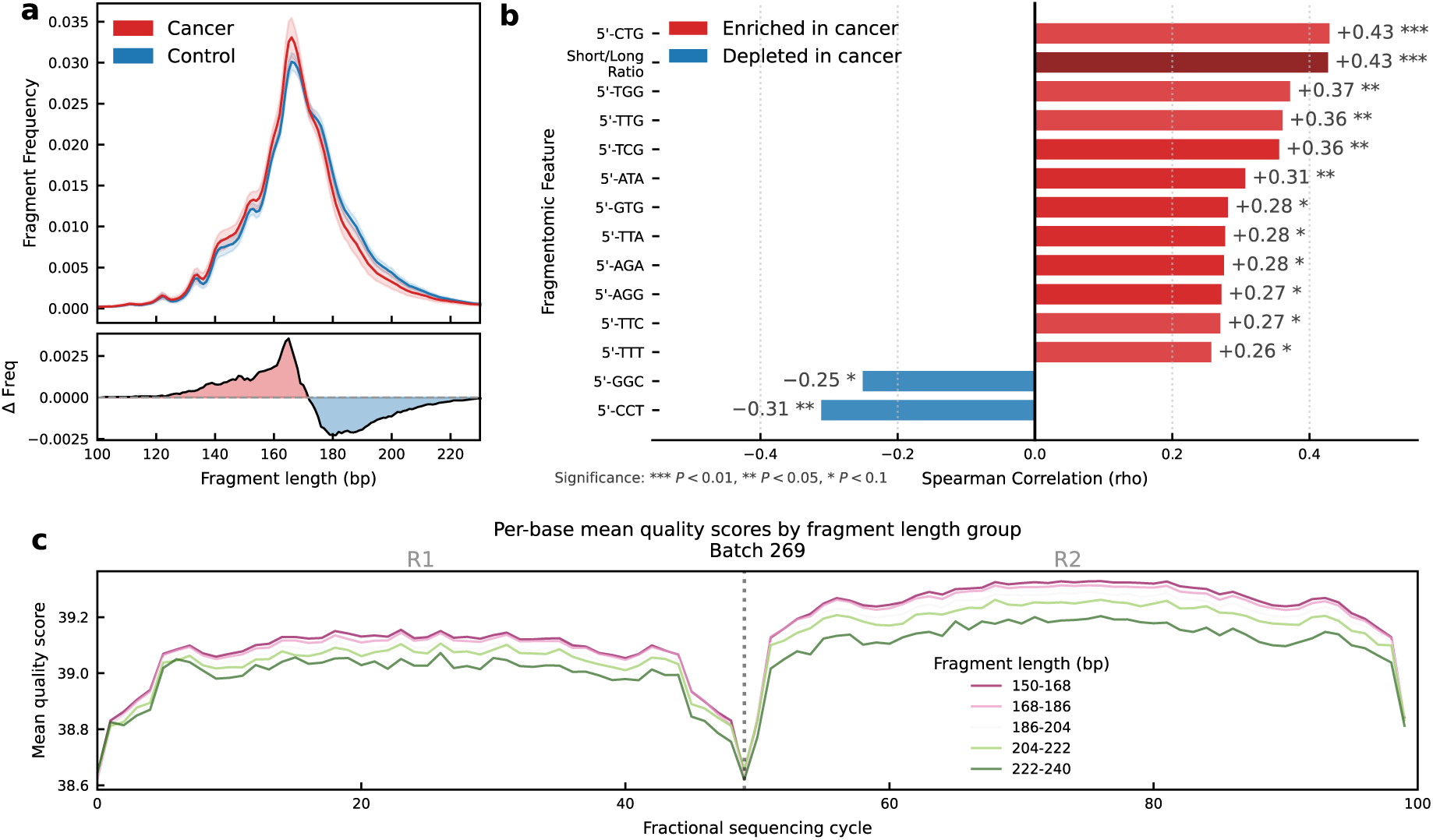
PBQS derived scores correlate with fragmentomic features. **a** Fragment size distribution on aggregated dataset. The top panel shows the frequency of fragment lengths for cancer (red) and control (blue) samples, revealing the characteristic enrichment of shorter cfDNA fragments in cancer. The bottom panel displays the differential frequency highlighting the specific enrichment of short fragments and depletion of longer fragments at the mononucleosomal peak (∼167 bp) **b** Spearman correlation analysis between the PC2 score and established fragmentomic features. Red bars indicate features known to be enriched in cancer (specific 5’ end-motifs and the short to long fragment ratio), while blue bars represent features depleted in cancer. The PC2 score exhibits significant positive correlations with cancer-associated markers and negative correlations with control-associated markers linking the quality score signal to underlying biological properties. Significance levels: *** *P* < 0.01, ** *P* < 0.05, * *P* < 0.1. **c** Impact of fragment length on per-base quality scores (Batch 269). Reads were stratified into length bins (colored lines) and their mean quality profiles were computed across the fractional sequencing cycle. Shorter fragments (purple) consistently exhibit higher mean quality scores compared to longer fragments (green), providing a mechanistic basis for how fragment length shifts in cancer modify the aggregate PBQS profile.

Notably, PC2 loadings and the ablation study results suggest that fragment ends are a significant factor driving condition separation. Several studies have reported systematic bias caused by distinct sequence composition^28,29^, such patterns may impair base-calling parameters, as phasing, when libraries have an unbalanced base composition in fragment ends. This observation aligns well with known end-motif characteristics of ctDNA and the utilization for cancer detection^30^.

To quantify the association between MFPQP and known fragmentomic features, we compared the PC2 scores with established fragmentomic biomarkers. We utilized the FrEIA framework (Moldovan *et al.*) to quantify all possible 5’ trinucleotide end-motifs and assess correlation with the ones found to be enriched and depleted in the pan-cancer study^31^. Additionally, we calculated overall fragment length ratio, defined as the ratio of short fragment (50-150 bp) to long fragments (250-1,000 bp). Spearman correlation analysis between these biological features and the PC2 derived score (Fig. 3b) suggest significant positive correlation with the short to long ratio (ρ = +0.43, *P* < 0.01) and with tumor-enriched end-motifs such as 5’-CTG (ρ = +0.43) and 5’-TGG (ρ = +0.37). Conversely, PC2 scores show significant negative correlations with healthy-associated motifs in control cfDNA such as 5’-CCT (ρ = −0.31).

Lastly, in the absolute-cycle PDAC subset, covariate models showed that cancer/control status remained associated with the absolute-cycle PBQS-derived score after adjustment for mean fragment length and mean fragment GC content, while the short-to-long fragment ratio was itself significantly associated with the score (Table S9). These results suggest that PBQS is related to fragmentation structure but is not reducible to mean fragment length or mean GC content alone.

These associations support a plausible suggestion of mechanism for why raw quality scores can be predictive: PC2 captures systematic, end-localized PBQS dynamics that are associated with fragment length and specific 5′ end-motifs. We therefore suggest that enrichment of shorter, motif-biased tumor-derived fragments can modestly modulate sequencing behavior near read termini, leaving a reproducible cancer-associated signature in the mean PBQS profiles.

### Comparison of PBQS-derived score with orthogonal methods

To evaluate agreement of our PBQS-derived score with orthogonal methods, we compared the out-of-fold PC2 values with three established ctDNA estimators, each focuses on a different ctDNA biomarker. Per-batch normalized FrEIA scores derived from 5’ trinucleotide end-motifs frequencies, mean DELFI scores computed from short to long fragment size ratios across genomic bins^23^, and ichorCNA, which calculates tumor burden estimates via copy number alterations analysis^32^ (Methods).

Correlation analysis across the entire dataset revealed a statistically significant positive association between PC2 scores and normalized FrEIA scores (Spearman ρ = 0.43, *P* = 0.003; Fig. 4a). This finding reinforces the hypothesis that the PBQS signal is driven by fragmentomic properties, specifically the overrepresentation of certain end-motifs in tumor-derived fragments that modulate sequencing qualities. Notably, this correlation was observed across both cancer and control groups, suggesting that PBQS captures the underlying biological gradient of fragment-end compositions.

**Fig. 4:**
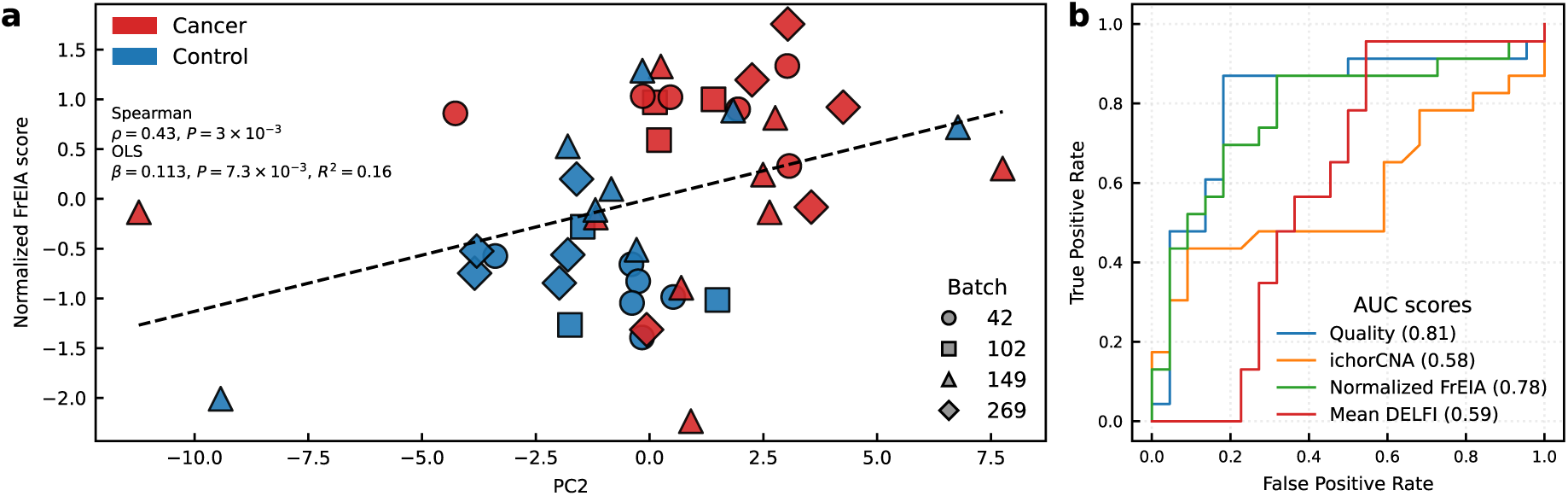
PBQS-derived PC2 scores as an orthogonal biomarker for cancer detection. **a** Correlation between PC2 and fragment-end motifs. Scatter plot illustrating the relationship between LOBO-projected PC2 scores and normalized FrEIA scores across the aggregated dataset. Points are colored by clinical label (red: cancer; blue: control) and shaped by sequencing batch. The significant positive correlation (Spearman ρ = 0.43, *P* = 0.003) suggests that the PBQS signal is a proxy for underlying biological fragment-end composition. **b** Comparative classification performance. ROC curves comparing the diagnostic accuracy of the PBQS-based PC2 classifier (AUC = 0.81) against established orthogonal methods: Normalized FrEIA (0.78), Mean DELFI (0.59), and ichorCNA (0.58). The PBQS-derived signal demonstrates superior or comparable performance to specialized fragmentomic and genomic pipelines in this cohort, despite relying solely on raw sequencing metadata.

Nevertheless, PC2 scores demonstrated minimal and nonsignificant correlation with fragmentation ratios as measured by the simplified mean DELFI calculation (Spearman ρ = 0.11, *P* = 0.44, Fig. S8a) and lack of association with tumor burden estimation from ichorCNA (Spearman ρ = −0.07, *P* = 0.62, Fig. S8b). The lack of correlation with ichorCNA is particularly noteworthy, as it suggests that the PBQS signal is orthogonal to large-scale genomic instability and remains detectable even in samples with low tumor purity where traditional CNV-based methods may struggle (Fig. S9).

Finally, a classification performance comparison using ROC analysis (Fig. 4b). In this specific cohort, the PBQS-based PC2 classifier achieved an AUC of 0.81, compared with ichorCNA (AUC = 0.58), mean DELFI (AUC = 0.59), and the motif-based FrEIA framework (AUC = 0.78). These results suggest that raw sequencing quality scores, which are typically treated as technical metadata, may capture fragmentomic information overlapping with motif-based methods such as FrEIA, while requiring substantially less computational overhead.

## Discussion

PBQS are typically treated as technical readouts used for read trimming and weighting evidence in variant calling. For the most part, quality variations are assumed to reflect sequencing chemistry, optics, or library preparation artifacts. Here, we demonstrate that in rigorously controlled cfDNA whole-genome sequencing batches, PBQS profiles also carry a subtle cancer-derived signal. This signal may represent underlying cfDNA fragmentomic properties rather than purely technical noise. Large case-control cfDNA sequencing datasets are available^23,33,34^, but they typically aggregate samples from multiple centers, flow-cell runs, and library-preparation kits. This introduces technical heterogeneity that confounds the analysis of PBQS profiles, making them inadequate for a proof-of-concept demonstration of biological manifestations. Thus, for PBQS-based analyses, dataset suitability depends not only on sample size, but also on whether cancer and control samples are matched within the same sequencing run or flow-cell context. To test our hypothesis, it was critical to eliminate technical contributors that could confound the analysis. This necessitated the use of a dataset where technical variables were minimized or controlled, ensuring that observed differences in quality scores could be more confidently attributed to biological factors. Factors such as the sequencing platform, library preparation protocols, and run parameters must be consistent across samples or appropriately accounted for.

In this study, we obtained and analyzed such a dataset. The assembled cohort is relatively small but rigorously controlled, representing patients and technically matched controls across several hospitals, cancer types, and processing regimes. All samples in each batch were handled and sequenced under a uniform protocol on a single flow-cell lane and further subsampled per tile. This approach substantially reduced technical confounders and allowed our analysis to focus on possible biological contributions to PBQS.

A key observation in the current work is that MFPQPs exhibit subtle but systematic differences between cancer and control samples. These differences are easily exposed via straightforward signal processing techniques, such as PCA. In both single-batch and aggregated analyses, the separation between cancer and controls is not driven by the dominant axis of variance (PC1), but instead by PC2, which captures boundary-enriched dynamics concentrated at the 5′ and 3′ fragment termini. This physical localization supports the interpretation that fragment-end composition and nucleosomal cleavage patterns, which are most apparent at fragment boundaries where end-motif features are expressed, contribute to the observed signal. The purpose of MFPQP is methodological: it enables comparison of batches sequenced with different read lengths. The persistence of the signal in absolute sequencing-cycle coordinates in the uniform 2×151 bp PDAC subset indicates that the observed separation is not solely an artifact of fractional-position mapping (Fig. S7).

Across four independent sequencing batches, the PC2-derived signal served as a consistent plane of separation. In batch-adjusted models, it maintained discriminative performance in a LOBO setting (macro-average AUC = 0.78 ± 0.15). Permutation tests, ablation results, and regression modeling further strengthen the conclusion that the observed LOBO performance is not driven by chance or technical noise; rather, the signal reflects structured differences between cancer and matched control cfDNA.

Our mechanistic interpretation suggests that PBQS partially encodes fragmentomic variation that modulates sequencing behavior near fragment boundaries. Fragment length stratification demonstrates a systematic relationship between fragment size and quality magnitude, with longer fragments showing lower mean PBQS across cycles. Additionally, PC2 correlates with established fragmentomic biomarkers, including a positive association with the short-to-long fragment ratio and tumor-enriched 5′ end-motifs (e.g., 5′-CTG and 5′-TGG), alongside a negative association with control-associated motifs (e.g., 5′-CCT). Additional covariate models further indicated that the PBQS-derived signal was not explained solely by mean fragment length or mean fragment GC content, although its association with the short-to-long fragment ratio supports a contribution from fragmentation structure (Table S9). These associations provide a plausible biological bridge: cancer-associated fragmentation skews cfDNA toward shorter, motif-biased fragments, which can influence SBS dynamics and leave a reproducible signature in aggregated PBQS profiles. Notably, while we identified these links, the influence of cancer-specific epigenetic modifications cannot be ruled out or confirmed by the current dataset and remains a potential avenue for future exploration.

While the quality-derived score shows significant correlation with motif-based FrEIA and fragment length ratios, it shows minimal association with simplified mean DELFI fragmentation scores and no association with ichorCNA tumor fraction estimates. This pattern suggests that PBQS may provide sufficient sensitivity for low-shedding, early-stage settings where genomic binning-based estimates are often weak.

More broadly, this work aligns with recent efforts to derive biologically informative phenotypes from complex biomedical data streams^35–37^. In this context, PBQS represents a routinely generated but underused sequencing-derived signal layer that may provide a lightweight surrogate of cfDNA fragmentomic properties.

From a practical standpoint, PBQS offers attractive operational advantages. Unlike other fragmentomic approaches that require full bioinformatic pipelines (including alignment, binning, and specialized feature extraction), PBQS is present in raw FASTQ files. It can be summarized immediately with minimal computational cost or the need for dedicated algorithms. This process yields a new type of biomarker for cfDNA analysis, enabling computationally lightweight screening as a standalone tool or as part of an ensemble with other fragmentomic and epigenomic features.

Nonetheless, several limitations temper the immediate clinical translation of this approach. The dataset presented here is small (*N* = 45); while intentionally designed to minimize technical confounders, the study lacks the statistical power to rigorously validate the hypothesis or stress-test the limit of detection. Future work should focus on validating the PBQS signature in larger, prospectively collected, multi-site, and multi-cancer cohorts with diverse pre-analytical conditions. Furthermore, robustness must be tested across various library processing kits and sequencing instruments. Because the current evidence remains associative, targeted mechanistic studies should also be performed to definitively link fragmentomics to quality dynamics. If generalizability is confirmed, PBQS could become a valuable, low-cost, and computationally efficient orthogonal biomarker in cfDNA diagnostics.

## Methods

### FASTQ normalization and downsampling

Raw FASTQ files were subjected to a custom preprocessing pipeline to ensure high-quality reads free of adapter contamination. Initially, adapters and indexes were trimmed using bbduk.sh (v. 39.22)^38^ in paired mode with the ktrim = r k = 23 mink = 11 hdist = 1 parameters and the adapter reference dataset provided with the software. Quality trimming was applied to the right end of the reads and a minimum length filter of 50 bp was enforced to discard fragments too short. Following trimming, fastp^7^ was utilized to generate quality control reports and validate the cleanliness of the processed data.

To mitigate potential batch effects arising from uneven sequencing coverage or flow cell edge effects, FASTQ files were normalized based on physical sequencing location. Read headers, adhering to the standard Illumina format, were parsed to extract coordinate information:

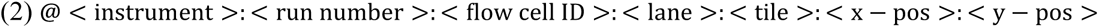

Subsequently, reads were parsed to extract ‘lane’ and ‘tile’ identifiers. Reads were grouped (binned) according to their lane and tile, and only bins providing at least 10,000 reads per sample were retained to ensure consistency among all samples. Within each retained bin, reads were randomly subsampled to exactly 10,000 per sample, resulting in uniform total read counts and balanced lane/tile representing an even number of reads for each sample per batch (Supplementary Table S1).

Fractional position quality profiles

To generate uniform quality metrics despite varying read lengths, per-base quality scores were mapped to a fixed-length vector of size *K* = 50. For a read of length *L* with quality scores *Q* = {*q*_0_, *q*_1_, …, *q*_*L*−1_}, the mapping of a base at position *i* to a bin *j* ∈ {0, …, *K* − 1} was calculated as:

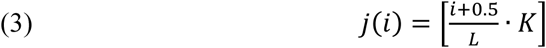

The aggregated quality score *S*_*j*_ for the *j*-th bin was computed as the mean quality of all bases falling into that bin:

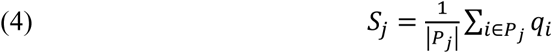

Where *P*_*j*_ = {*i* ∣ *j*(*i*) = *j*} is the set of positions in the read mapping to bin *j*. This process yielded a standardized 50 × 1 quality vector per read, from which per base fractional quality mean vector per sample was calculated. Fractional mean quality scores from R1 and R2 were concatenated resulting in a 100 vector per sample for the downstream analysis. To account for technical variation arising from different sequencing runs, batches, fractional quality vectors were standardized using Z-score normalization independently within each batch.

### Statistical analysis and dimensionality reduction

A LOBO cross-validation framework was employed to assess the robustness of the biological signal. In each iteration, PCA was trained on data from all batches except one and the held-out batch was projected into the learned latent space. PC2 was evaluated as the primary axis separating cancer and control samples. To ensure directional consistency across folds, the sign of the projected PC2 scores was aligned such that positive values corresponded to the cancer class based on the training set means. Model performance was evaluated using the AUC, reported as both a global pooled AUC (calculated on the concatenated out-of-fold predictions) and a macro-average AUC across batches.

Statistical validation was performed to confirm that the observed signal was not driven by batch effects. An OLS regression model (*PC*2 ∼ *Label* + *Batch*) was fitted to the standardized PC2 scores to test the significance of the cancer/control label while controlling for batch identity. Additionally, effect sizes were quantified using Cohen’s *d* for PC1 and PC2 within each batch, summarized via sample-size weighted averaging. Finally, statistical significance was determined using a stratified permutation test (10,000 iterations), where labels were shuffled randomly within their respective batches to generate a null distribution of LOBO performance metrics, preserving the original batch structure and class imbalance.

Additional OLS covariate models were used to assess whether the PBQS-derived signal was explained by fragment-length or GC-content summaries. For the PDAC absolute-cycle subset batches, the LOBO PC2 score was modeled as a function of cancer/control label, sequencing batch, mean fragment length, mean fragment GC content, and short-to-long fragment ratio.

### Absolute-cycle analysis controls for fractional-position encoding

As a control for the fractional-position representation, we repeated the PBQS analysis in absolute sequencing-cycle coordinates using only the three PDAC batches sequenced under a uniform 2×151 bp protocol. For each sample, mean PBQS values were computed for cycles 1–151 of R1 and cycles 1– 151 of R2 and concatenated into a 302-dimensional vector. These absolute-cycle vectors were batch-standardized and analyzed using the same PCA and LOBO framework.

### Data processing for orthogonal validation

For scoring samples using FrEIA, DELFI and ichorCNA each sample was aligned to the reference genome using the nf-core Sarek pipeline^39^ with BWA-MEM^40^ as the alignment tool. Analysis of ichorCNA was performed using ichorCNA (v. 0.5.0.) All aligned sequencing data in BAM format, including clinical samples and matched technical controls, underwent standardized pre-processing pipeline to generate corrected read-depth profiles for copy number analysis. This procedure was implemented using the HMMcopy_utils^41^ suite. Genome-wide read counts were quantified by binning the genome into non-overlapping windows of a fixed size. The readCounter utility was employed for this task. The following changes to default ichorCNA parameters were used:

centromere=’GRCh38.GCA_000001405.2_centromere_acen.txt’, scStates=’’, repTimeWig=’’, ploidy=2, includeHOMD=FALSE, estimateNormal=TRUE, estimatePloidy=TRUE, altFracThreshold=0.05, genomeBuild=“hg38”, genomeStyle=“UCSC”, flankLength=100000, txnE=0.9999999, chrs=“c(’chr1’,…,’chr22’)”, chrTrain=“c(’chr1’,…,’chr22’)”, maxCN=7, estimateScPrevalence=TRUE, normal=“c(0.2,0.3,0.4,0.5,0.6,0.7,0.8,0.9,0.95,0.97,0.98,0.99)”, initial_ploidies=c(2, 3, 4), maxFracGenomeSubclone=0.5, maxFracCNASubclone=0.7

A Panel of Normals (ichorCNA parameter “normal_panel”), mappability and GC files for 500kb were also taken from the ichorCNA resource. https://github.com/broadinstitute/ichorCNA

FrEIA (Fragment End Integrated Analysis) was employed to extract the frequencies of 5’ fragment end sequences. A normalized FrEIA score was calculated as the ratio of the summed frequencies of the enriched and depleted set of 16 tumor-associated trinucleotides (AAG, ACG, AGA, AGG, ATA, ATG, CCG, GAG, GCG, GGG, GTA, GTG, TCG, TTA, TTC, TTG) to a set of 15 control-associated trinucleotides (ACA, ACC, ACT, CAC, CAT, CCA, CCC, CCT, CGA, CGC, CGT, CTT, GAC, GGC,

TCT) described by Moldovan *et al.,* To mitigate technical batch effects, the resulting ratios were standardized (Z-score normalized) within each processing batch prior to downstream comparison. https://github.com/mouliere-lab/FrEIA

Mean DELFI (DNA EvaLuation of Fragments for Early Interception) scores were calculated per sample using the FinaleToolkit^42^ (v. 0.11.0) and the delfi command while using default suggested parameters and computing the mean across the corrected fragment length ratios. https://github.com/epifluidlab/FinaleToolkit

### Patient recruitment and sample collection

This study was conducted in accordance with the ethical standards of the institutional research committees. The study was approved by the Helsinki Committee of the Sheba Medical Center (approval number 9534-22-SMC) and the Helsinki Committee of the Tel Aviv Sourasky Medical Center (approval number 0569-16-TLV). Informed consent was obtained from all participants prior to sample collection.

The PDAC cohort included patients at stages I to IV, with samples collected at various clinical time points (e.g., pre-adjuvant, on-treatment, post-resection). The breast cancer cohort consisted of female patients diagnosed predominantly with infiltrating ductal carcinoma (Stages I-II, based on pTNM staging) with varying hormone receptor statuses (ER, PR, Her-2). The control group for both cohorts included individuals with no known malignancy or, for the breast cancer cohort, individuals with benign conditions such as fibroadenoma.

Across both cohorts, participants ranged in age from 21 to 80 years. Blood was collected using ethylenediaminetetraacetic acid (EDTA) tubes for all participants, except for one healthy control sample collected in a Streck tube.

### Plasma separation and cfDNA extraction

For the PDAC cohort, whole blood samples were processed promptly in-house to isolate plasma. The procedure involved an initial centrifugation at 1,500 × g for 10 minutes at 4°C using Lymphoprep™ density gradient medium, followed by a second centrifugation of the separated plasma at 3,000 × g for 10 minutes at 4°C to remove residual cellular debris. For the breast cancer cohort, plasma was separated at the collection site prior to its arrival at our lab. For all samples, the final clarified plasma was aliquoted and stored at –80°C until cfDNA extraction.

The plasma from patients with breast cancer and benign tumors were obtained from the TLV BioBank, Tel Aviv Sourasky Medical Center, Tel Aviv, Israel. For this cohort, whole blood samples were collected in EDTA tubes, centrifuged at 1,200 × g for 15 minutes at room temperature, and the resulting plasma was aliquoted and stored at –80°C until cfDNA extraction. Subsequently, cfDNA was extracted using the QIAamp® Circulating Nucleic Acid Kit (Qiagen), following the manufacturer’s protocol. The starting plasma volume varied between cohorts, with 2–4 mL used for the PDAC samples and 0.5 mL for the breast cancer samples. Eluted cfDNA was quantified using the Qubit™ 4 Fluorometer with the dsDNA High Sensitivity (HS) Assay Kit. The differences in plasma volume input were reflected in the final yields; cfDNA concentrations ranged from 0.13 to 2.44 ng/μL across both cohorts, yielding total amounts between 3 and 15 ng per sample.

### cfDNA quality assessment

The quality and fragment size distribution of the extracted cfDNA were evaluated using the Agilent 4200 TapeStation System with the Cell-free DNA ScreenTape assay. This platform allows for the determination of the proportion of cfDNA within the expected size range for mononucleosomal DNA. The quality metrics varied between the cohorts. Samples from the PDAC cohort displayed a high degree of purity and fragment integrity, with cfDNA percentages ranging from 77% to 90%. In contrast, the breast cancer cohort showed a wider and lower range of cfDNA percentages (35% to 73%). Although some samples fell below the typical quality control threshold of 70%, they were retained for the study as they successfully produced high-quality sequencing data, confirming their suitability for downstream analysis.

### Library preparation and whole-genome sequencing

Sequencing libraries were prepared and sequenced for the two clinical cohorts at different facilities using distinct protocols. All resulting raw sequencing data in FASTQ format were then processed through the same standardized alignment and pre-processing pipeline^43^.

For the breast cancer cohort (*n* = 19), following DNA extraction, libraries were generated using a standard ligation-based workflow for whole-genome resequencing. The prepared libraries were then sequenced on an Illumina platform to generate 2×143 bp paired end reads. The sequencing run yielded a mean of 22.1 Gb of data per sample.

For the PDAC cohort (*n* = 21), sequencing libraries were prepared from cfDNA using the Accel-NGS 2S PCR-Free kit, which is optimized for low-input samples and minimizes amplification bias. The libraries were subsequently sequenced on an Illumina NovaSeq X Plus platform to generate 2×151 bp paired end reads. This process targeted an output of 10 Gb of data per sample.

## Data availability

The raw sequencing data generated in this study has been deposited in the European Genome-phenome Archive (EGA) under the study accession number EGAS50000001620. Processed sequence data needed to reproduce the analyses demonstrated here are available for download on Zenodo at https://doi.org/10.5281/zenodo.18909774.

## Contributions

H.V. designed, planned and executed the study. M.G. and R.S. performed the biological experiments. M.R.G., T.G and T.R. assisted in sample acquisition. D.L. provided statistical assistance and N.S. provided overall scientific supervision, critically revised the manuscript, read and approved the final version.

## Acknowledgements

We thank members of the Shomron laboratory for handling the biological processing and DNA extraction along valuable support in data acquisition and technical consultation. We thank Dr. Moni Shahar from the AI and Data Science Center, Prof. Asaf Madi from the School of Medicine, Prof. Yuval Ebenstein from the School of Chemistry, Tel Aviv University, Israel, and Prof. Yaniv Erlich for their willingness to read, share thoughts and engage in fruitful discussion. We thank the TLV BioBank, Tel Aviv Sourasky Medical Center, Tel Aviv, Israel, for providing the breast cancer and control patients cohort.

---

**Fig. S1:**
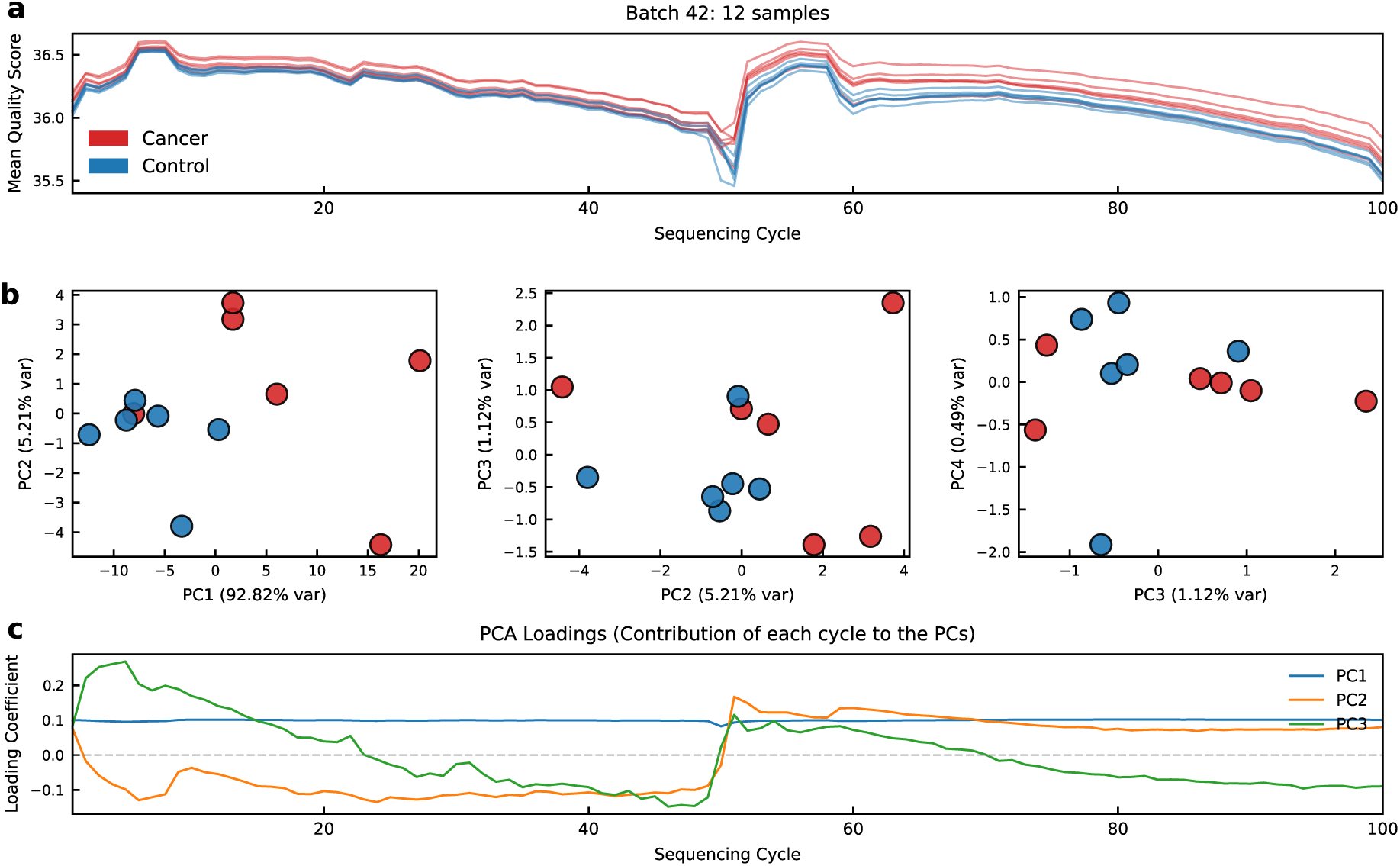
Unsupervised analysis of PBQS for Batch 42. **a** MFPQP for Batch 42 (*n* = 12). The traces represent the aggregated per-base quality scores mapped to a standardized fractional vector (*K* = 100) for PDAC patients (red) and healthy controls (blue). **b** PCA projections. The panels display the relationships between the first four principal components (PC1 vs. PC2, PC2 vs. PC3, and PC3 vs. PC4). Despite the smaller sample size, separation between cancer and control is observable along the PC2 axis. **c** PCA Loadings vectors. The coefficients indicate the contribution of each fractional sequencing cycle to the first three principal components. Consistent with the main cohort, PC2 (orange) exhibits dynamic variance at the read boundaries.

**Fig. S2:**
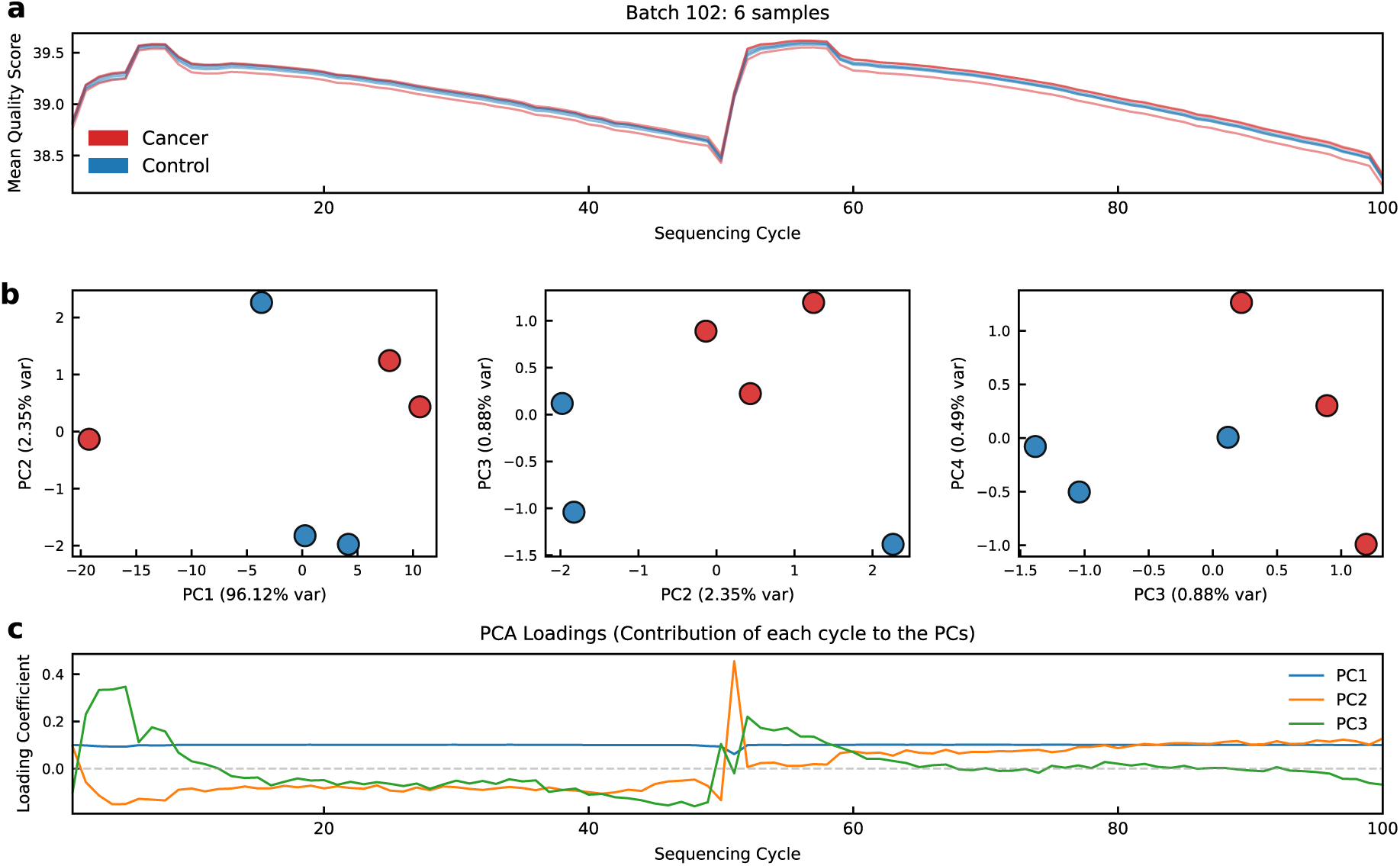
Unsupervised analysis of PBQS for Batch 102. **a** MFPQP for Batch 102 (*n* = 6). Traces show the standardized quality profiles for the smallest PDAC cohort in the study. **b** PCA projections. Scatter plots illustrate the distribution of samples in the latent space defined by the first four principal components. **c** PCA Loadings vectors. The loading profiles for PC1 (blue), PC2 (orange), and PC3 (green) confirm that the boundary-driven signal (PC2) persists even in this limited sample set, distinct from the global mean intensity tracked by PC1.

**Fig. S3:**
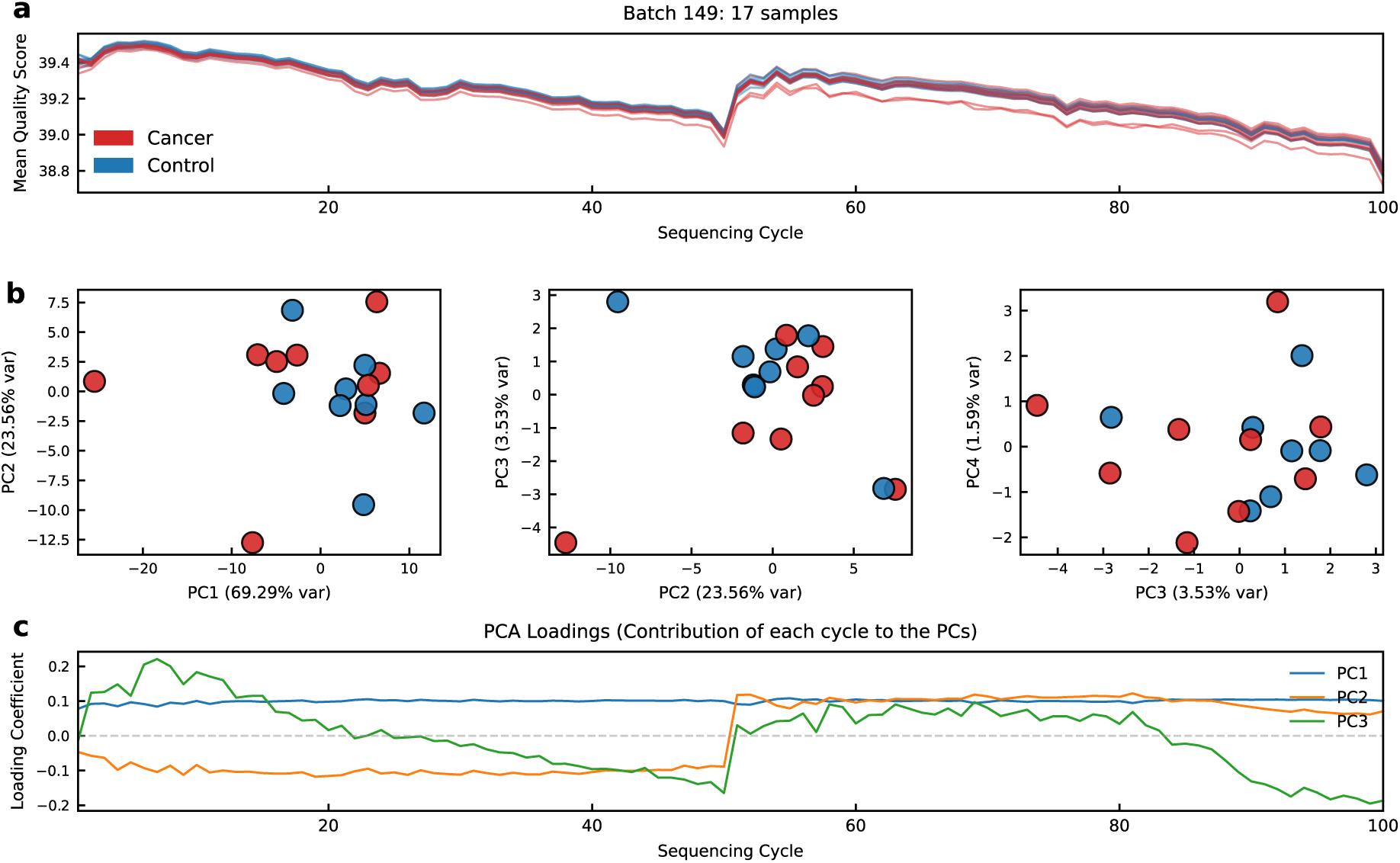
Unsupervised analysis of PBQS for Batch 149. **a** MFPQP for Batch 149 (*n* = 17). This batch represents the breast cancer cohort (red) and benign controls (blue) sequenced with a different read length (143 bp). **b** PCA projections. The plots display the separation of samples using the first four principal components. This batch validates the transferability of the signal to a different cancer type and sequencing regime. **c** PCA Loadings vectors. The loading coefficients demonstrate that the “wavy” boundary signal in PC2 (orange) is a consistent feature of cancer-derived cfDNA, independent of the specific sequencing platform or cancer indication.

**Fig. S4:**
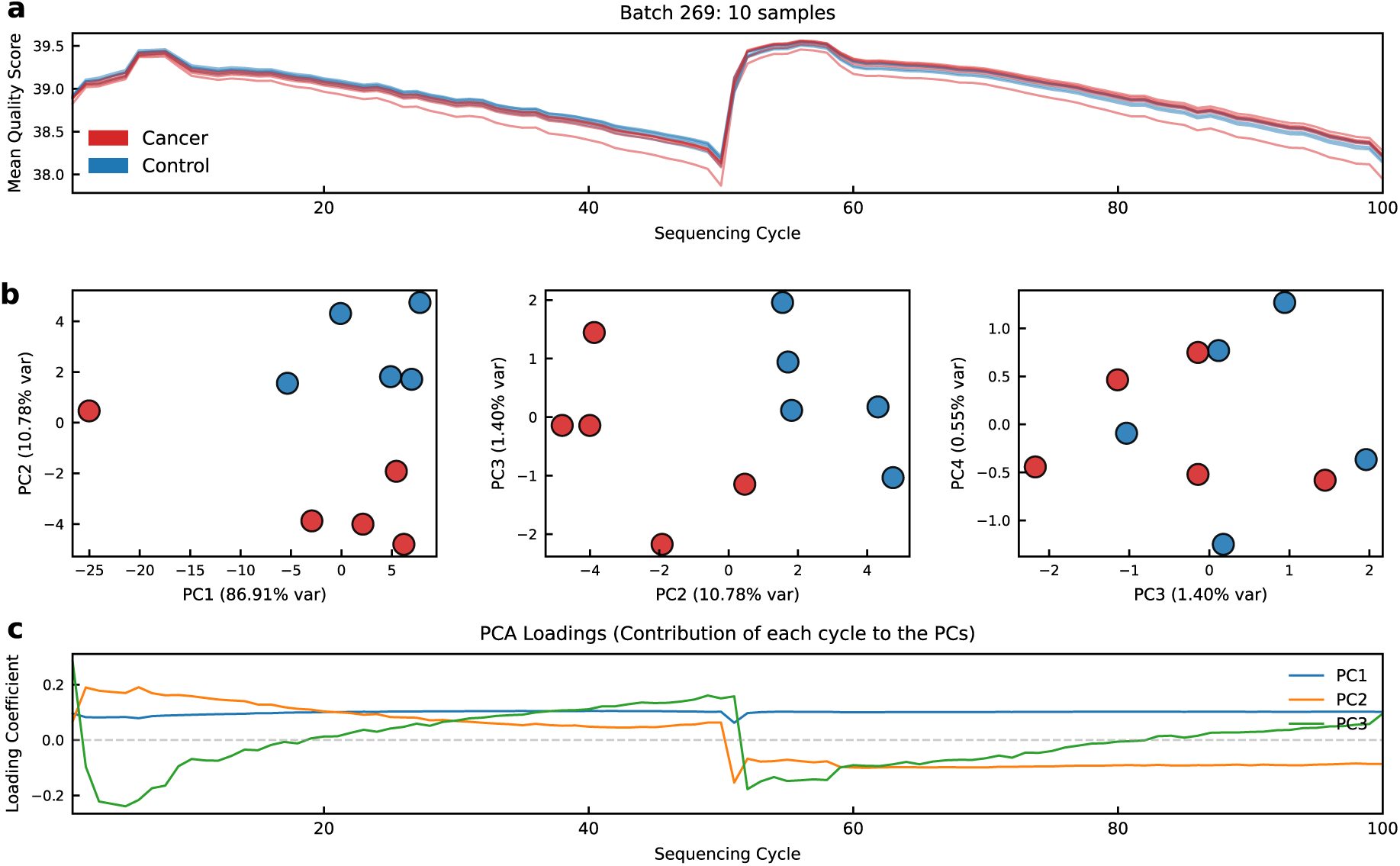
Unsupervised analysis of PBQS for Batch 269. **a** MFPQP for Batch 269 (*n* = 10). Traces show the standardized quality profiles for the representative PDAC cohort from Fig. 1. **b** PCA projections. Scatter plots illustrate the distribution of samples in the latent space defined by the first four principal components. **c** PCA Loadings vectors. The loading profiles for PC1 (blue), PC2 (orange), and PC3 (green) confirm that the boundary-driven signal (PC2).

**Fig. S5:**
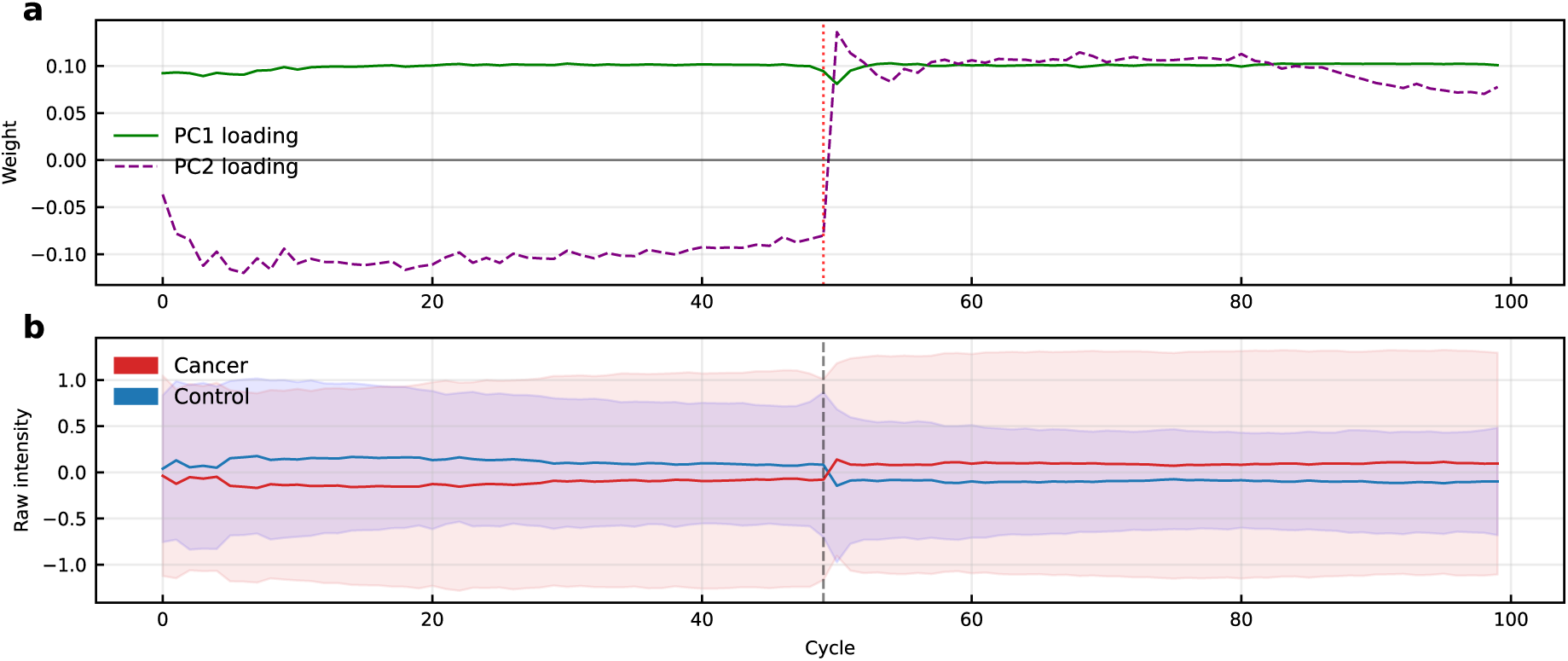
Physical interpretation of the first two PCs. **a** Visualization of the PCA loading vectors across the full dataset. PC1 loading vector (green) remains flat and positive, suggesting that it tracks the global mean quality magnitude of the samples. In contrast, PC2 loading vector (purple dashed) exhibits distinct dynamic fluctuations, particularly at the 5’ start sites of both reads (indices 0–10 and 50–60). The vertical red dotted line marks the concatenation boundary between paired end reads. **b** Aggregated mean quality profiles for the entire dataset. The solid lines display the average signal intensity for cancer (red) and control (blue) samples across the mean fractional sequencing cycle, with shaded regions representing the standard deviation. The vertical grey dashed line indicates the junction between R1 and R2. The systematic separation between the groups corresponds to the specific profile dynamics captured by the PC2 weights shown in **a**.

**Fig. S6:**
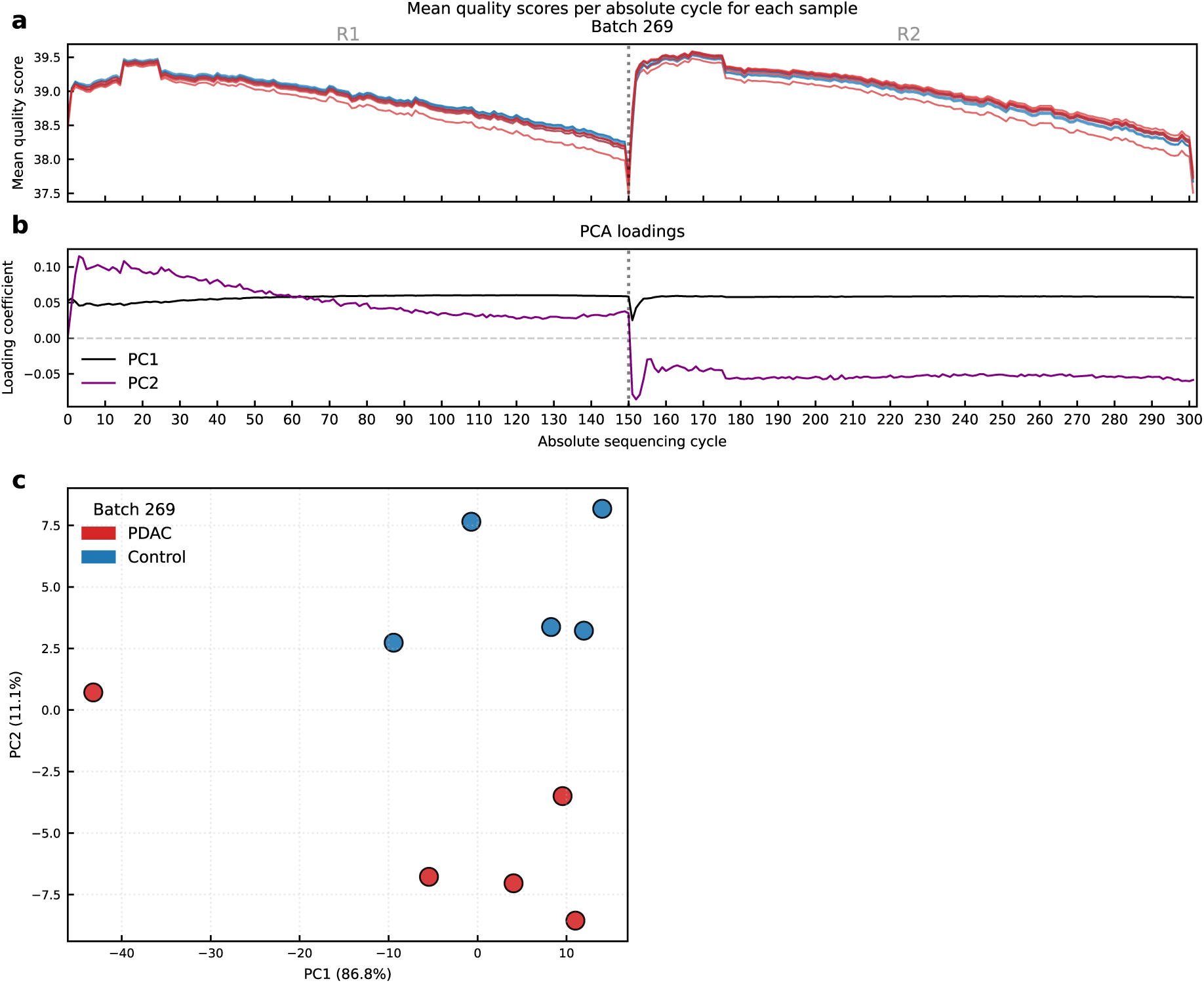
Absolute-cycle PBQS analysis of representative PDAC batch 269. **a** Absolute cycle for Batch 269. The traces display the aggregated per-base quality scores mapped to the absolute cycle vector (302, concatenating Read 1 and Read 2). Cancer samples exhibit a subtle but systematic elevationA in quality scores compared to controls. **b** PCA loading vectors for the first two principal components. The PC1 loading vector (black) remains relatively flat across the sequencing cycle. In contrast, the PC2 loading vector (purple). **c** PCA projection of a representative PDAC batch (Batch 269). The scatter plot reveals a distinct separation between cancer samples (red) and controls (blue) along the second principal component (PC2), despite PC1 capturing the majority of the variance.

**Fig. S7:**
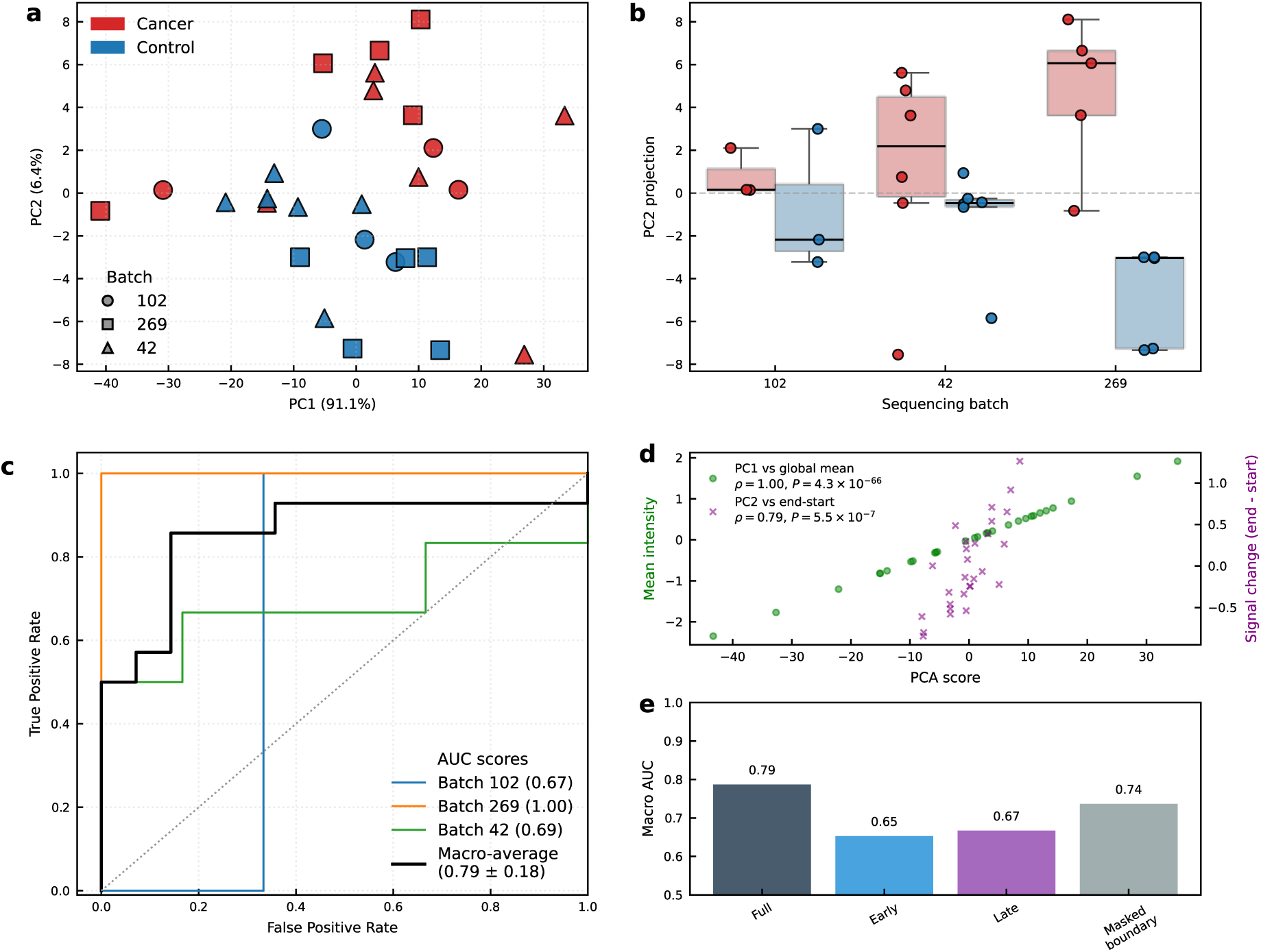
Absolute-cycle PBQS analysis across the three 2×151 bp PDAC batches. **a** PCA projection of the aggregated per batch Z-scored 2×151 bp batches only shaped by sequencing batch (red: cancer, blue: control). While PC1 captures the majority of variance (91.1%), PC2 (6.4%) defines the axis of separation between cancer and control samples. **b** Distribution of PC2 scores stratified by sequencing batch. In all three batches, cancer samples exhibit consistently higher PC2 scores compared to their matched technical controls. **c** ROC curves from LOBO cross-validation. Colored lines indicate performance when testing on each specific held-out batch, while the black line represents the macro-average performance (AUC = 0.79 ± 0.18) **d** Correlation analysis linking latent PCA components to physical quality attributes. PC1 (green circles) shows perfect correlation with the global mean quality score (Pearson’s ρ = 1.00). PC2 (purple crosses) correlates significantly with the “end-start” quality delta (Pearson’s ρ = 0.79). **e** Feature ablation study comparing LOBO performance across different segments of the quality profile. The “Masked boundary” model, preserves classification power (AUC = 0.74), whereas models restricted to only the early or late segments perform at or below random chance.

**Fig. S8:**
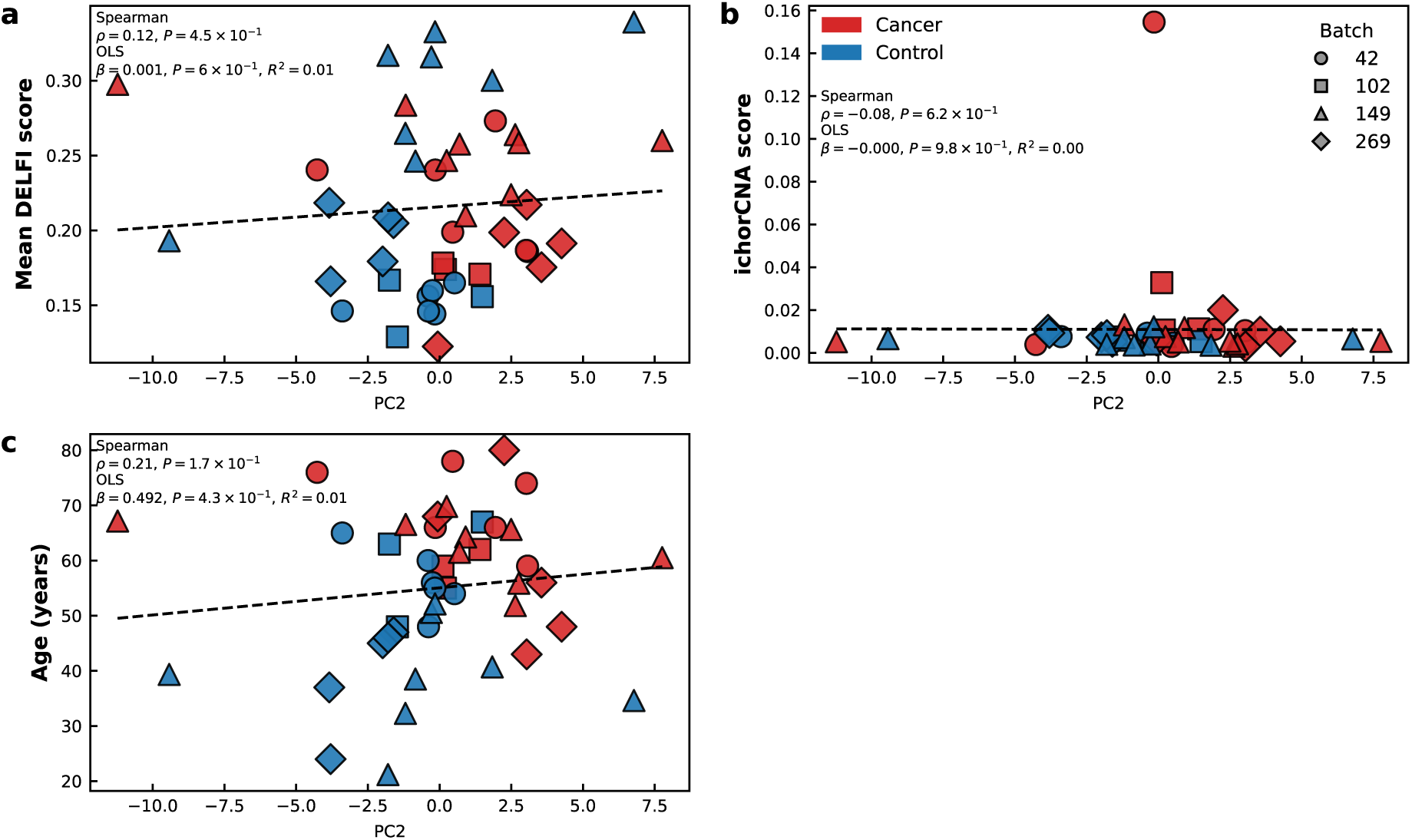
Orthogonality of PBQS-derived signal to fragment size ratios, copy number alterations and age. **a** Correlation between PC2 and Mean DELFI scores. Scatter plot showing the relationship between out-of-fold PC2 scores and the mean DELFI fragmentation score. Points are colored by clinical label (red: cancer; blue: control) and shaped by sequencing batch. Statistical analysis reveals no significant correlation (Spearman ρ = 0.12, *P* = 0.45), indicating that the PBQS signal captures biological information distinct from broad fragment size distributions. **b** Correlation between PC2 and ichorCNA scores. Scatter plot comparing PC2 scores with tumor fraction estimates derived from copy number analysis. No significant association was observed (Spearman ρ = −0.08, *P* = 0.62), suggesting the PBQS-derived biomarker is independent of large-scale genomic instability. **c** Correlation between PC2 and patient age. Scatter plot evaluating age as a potential demographic confounder. No significant global correlation was observed (Spearman ρ = 0.21, *P* = 0.17; OLS *R*^2^ = 0.01, *P* = 0.43), demonstrating that the malignancy-associated PBQS signature is not a byproduct of healthy aging processes and remains robust across the entire age spectrum (21–80 years). In all panels, the dashed line represents the OLS regression fit, and the flat trajectories across all metrics underscore the orthogonality of the PBQS-derived PC2 marker to standard clinical and fragmentomic confounders.

**Fig. S9:**
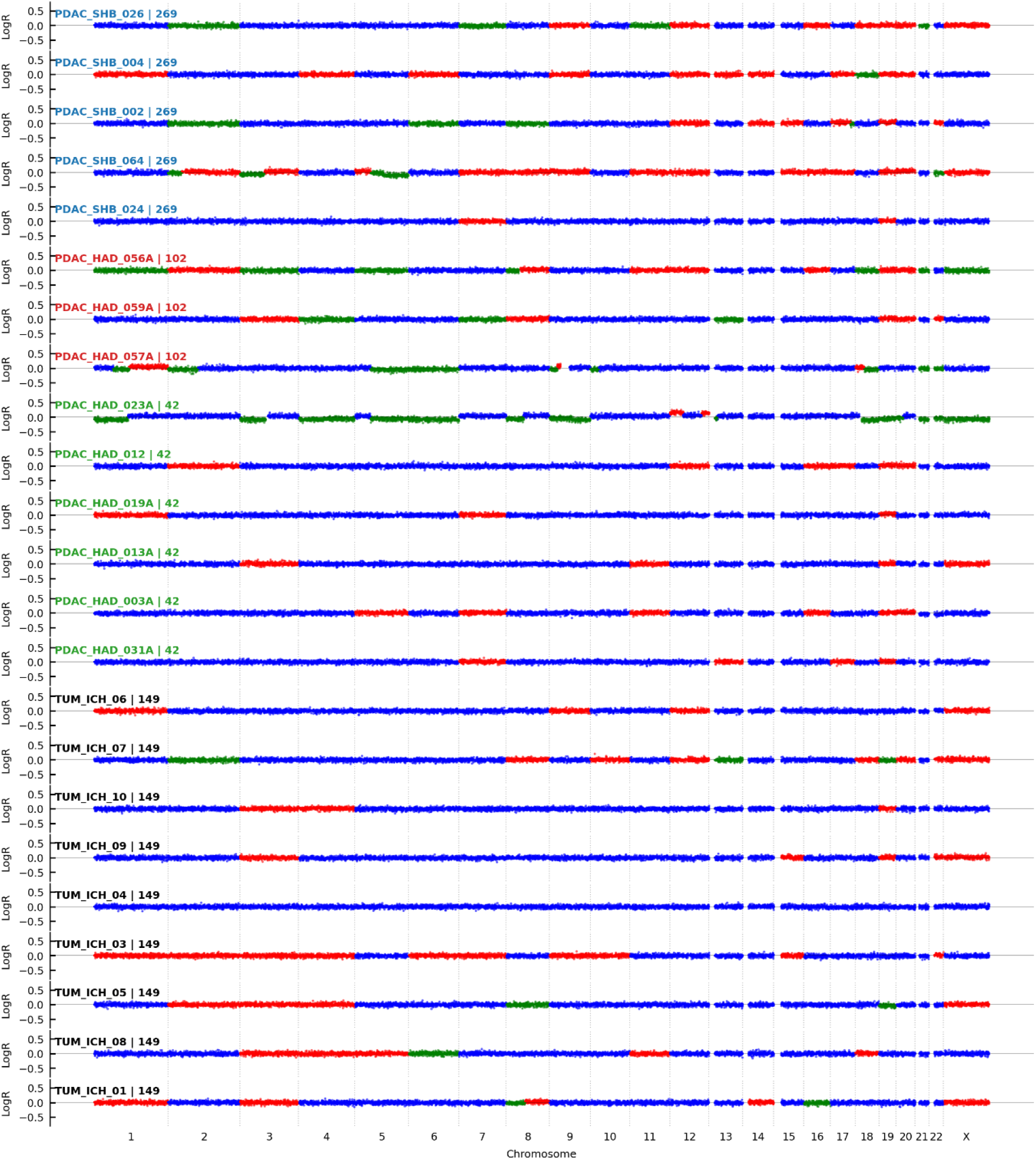
Genome-wide copy number profiles for cancer samples Individual genome-wide copy number profiles (LogR) generated by ichorCNA for cancer samples across all four sequencing batches (269, 102, 42, and 149). Each panel displays the log2-ratio of read depth across chromosomes 1–22 and X. The predominantly flat profiles observed across the majority of samples indicate a lack of large-scale, detectable copy number alterations (CNAs). This visualization underscores the relatively low tumor fraction or genomic stability in these clinical samples, highlighting the efficacy of the PBQS-based PC2 score in detecting cancer-specific signals even where traditional CNV-based detection methods show limited signal.

**Table S1.**
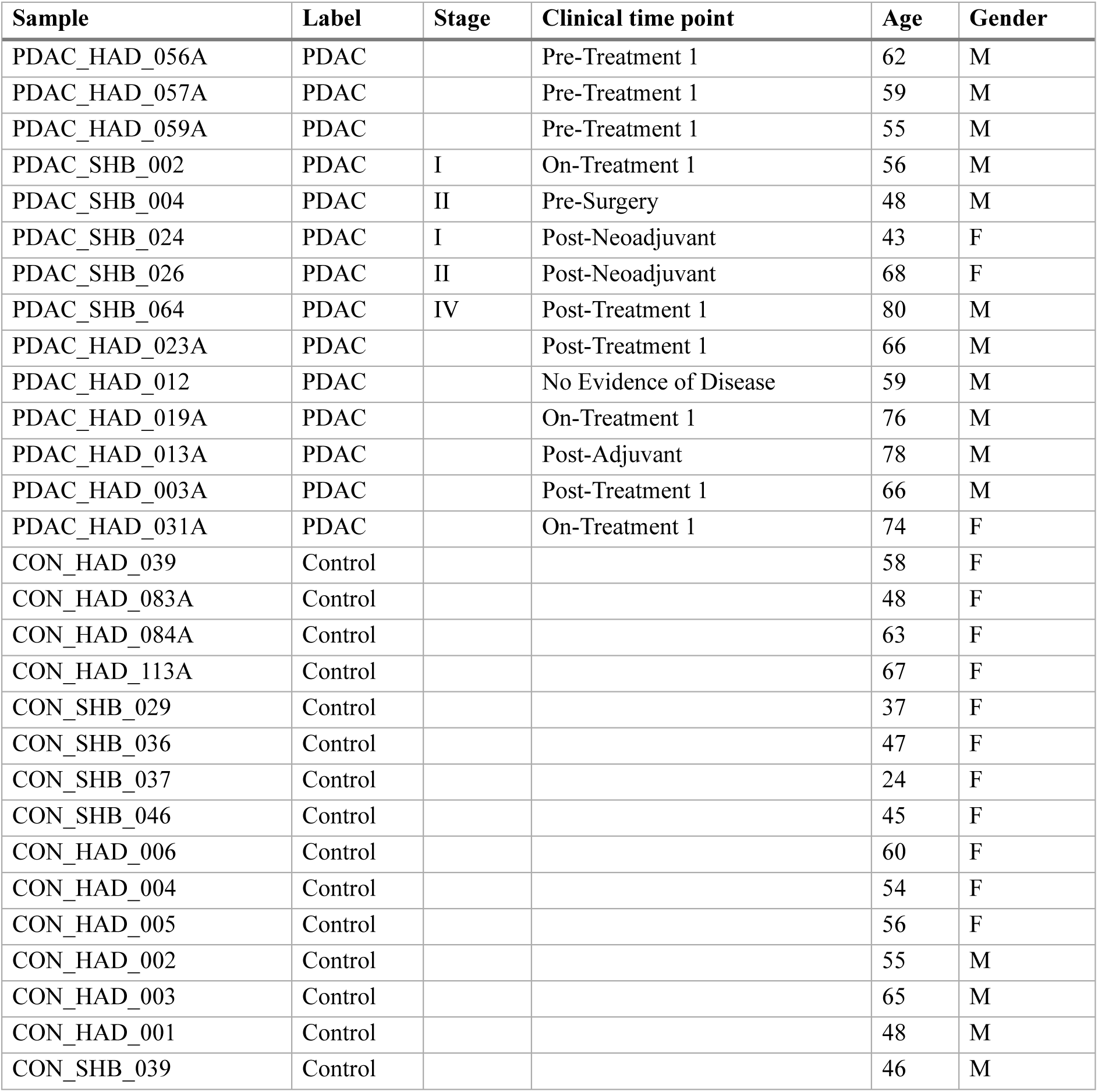
Demographic and clinical characteristics of the PDAC samples. Patient metadata at clinical time point for PDAC and matched control samples.

**Table S2.** Demographic and clinical characteristics of the breast cancer samples. Pathological pTNM classifications describe the extent of disease: pT denotes primary tumor size (pT1: ≤ 2 cm; pT2: 2–5 cm) and pN denotes regional lymph node involvement (N0: no metastasis; N1/N1a: 1–3 local nodes; N1(mi): micrometastases 0.2–2.0 mm; N2: 4–9 nodes; Nx: status unknown). This cohort is comprised of localized, stage I-II patients. Control samples represent biopsy-confirmed benign findings.

| Sample | Label | Finding Nature | Grade | Gender | Age |
| --- | --- | --- | --- | --- | --- |
| TUM_ICH_01 | Cancer | Infiltrating ductal carcinoma | pT1N2 | F | 61.5 |
| TUM_ICH_03 | Cancer | Infiltrating ductal carcinoma | pT2N1 | F | 67.17 |
| TUM_ICH_04 | Cancer | Infiltrating ductal carcinoma | pT1N1a | F | 51.83 |
| TUM_ICH_05 | Cancer | Infiltrating ductal carcinoma | pT1 N1(mi) | F | 55.88 |
| TUM_ICH_06 | Cancer | Infiltrating ductal carcinoma | pT2N0 | F | 60.53 |
| TUM_ICH_07 | Cancer | Infiltrating ductal carcinoma | pT2N0 | F | 64.27 |
| TUM_ICH_08 | Cancer | infiltrating ductal carcinoma | pT2N0 | F | 65.64 |
| TUM_ICH_10 | Cancer | infiltrating ductal carcinoma | pT2Nx | F | 69.82 |
| TUM_ICH_09 | Cancer | Infiltrating ductal carcinoma | pT2N0 | F | 66.55 |
| CON_ICH_01 | Control | Fibroadenoma |  | F | 21.18 |
| CON_ICH_02 | Control | fibrotic stroma with adenosis, fibrocystic and lactational changes |  | F | 34.62 |
| CON_ICH_03 | Control | Fibroadenoma |  | F | 32.29 |
| CON_ICH_05 | Control | Fibroadenoma |  | F | 50.58 |
| CON_ICH_06 | Control | fibroadenoma |  | F | 38.51 |
| CON_ICH_07 | Control | Fibroadenoma |  | F | 52.14 |
| CON_ICH_08 | Control | fibroadenoma |  | F | 40.72 |
| CON_ICH_09 | Control | Fibroadenoma |  | F | 39.35 |

**Table S3.** Batch effect analysis (Cohen’s *d*) Effect size of the separation between cancer and control samples for PC1 and PC2 within each batch.

| Batch | N (cancer/control) | PC1 Cohen's *d* | PC2 Cohen's *d* |
| --- | --- | --- | --- |
| 102 | 6 (3/3) | -0.040 | 0.843 |
| 149 | 17 (9/8) | -0.696 | 0.250 |
| 269 | 10 (5/5) | -0.578 | 3.678 |
| 42 | 12 (6/6) | 1.571 | 0.624 |
| Weighted Average | 45 | 0.022 | 1.190 |

**Table S4.** OLS regression model summary.

| Variable | Coefficient | Std. error | t-statistic | P-value |
| --- | --- | --- | --- | --- |
| Intercept | -0.3974 | 0.423 | -0.939 | 0.353 |
| Label [cancer] | 0.7947 | 0.290 | 2.736 | 0.009 |
| Batch [cancer.149] | -0.0234 | 0.462 | -0.051 | 0.960 |
| Batch [cancer.269] | -1.35e-16 | 0.503 | -0.000 | 1.000 |
| Batch [cancer.42] | 1.37e-16 | 0.487 | 0.000 | 1.000 |

**Table S5.** LOBO cross-validation results. Performance of the PC2-based classifier when testing on unseen batches.

| Test batch | Cancer | N test samples | AUC |
| --- | --- | --- | --- |
| 102 | PDAC | 6 | 0.67 |
| 149 | Breast | 17 | 0.69 |
| 269 | PDAC | 10 | 1.00 |
| 42 | PDAC | 12 | 0.78 |
| Global Pooled | All | 45 | 0.81 |
| Macro Average | All | - | 0.78 ± 0.15 |

**Table S6.** Feature ablation study. Macro-average AUC performance using subsets of the quality profile.

| Feature Set | Description | Macro AUC |
| --- | --- | --- |
| Full Series | All 100 bins | 0.78 |
| Masked Boundary | First 30% + Last 30% bins | 0.76 |
| Late Only | Last 33% bins | 0.42 |
| Early Only | First 33% bins | 0.27 |

**Table S7:** Initial-cycle masking and absolute 10 bp bin contribution analyses. Classification performance after removing the first 3, 5, or 10 bases from both reads, and predictive contribution of fixed 10 bp absolute-cycle bins in the PDAC 2×151 bp subset. Direction-corrected AUC was calculated as max(AUC, 1−AUC) to quantify bin-level separability independent of sign. The PBQS signal remained stable after initial-cycle masking, and informative 10 bp bins were distributed beyond the first 10 bases of either read.

| Feature set | First bases removed from each read | Features used | Pooled AUC | Macro AUC ± SD | Mann–Whitney *P* | Effect size |
| --- | --- | --- | --- | --- | --- | --- |
| All cycles | 0 | 302 | 0.857 | 0.787± 0.185 | 0.0014 | 1.114 |
| Remove first 3 bp | 3 | 296 | 0.837 | 0.787± 0.185 | 0.0026 | 1.101 |
| Remove first 5 bp | 5 | 292 | 0.847 | 0.787± 0.185 | 0.0019 | 1.107 |
| Remove first 10 b | 10 | 282 | 0.862 | 0.796± 0.179 | 0.0012 | 1.136 |

| Read | Absolute cycles | Pooled AUC | Direction-corrected AUC | Macro AUC ± SD | Mann–Whitney *P* | Effect size |
| --- | --- | --- | --- | --- | --- | --- |
| R1 | 1–10 | 0.357 | 0.643 | 0.304 ± 0.119 | 0.206 | -0.412 |
| R1 | 11–20 | 0.77 | 0.77 | 0.834 ± 0.221 | 0.0159 | 1.018 |
| R1 | 21–30 | 0.214 | 0.786 | 0.200 ± 0.077 | 0.0108 | -0.931 |
| R1 | 31–40 | 0.648 | 0.648 | 0.671 ± 0.307 | 0.19 | 0.592 |
| R1 | 41–50 | 0.245 | 0.755 | 0.302 ± 0.229 | 0.0229 | -0.827 |
| R1 | 51–60 | 0.49 | 0.51 | 0.566 ± 0.382 | 0.945 | 0.027 |
| R1 | 61–70 | 0.372 | 0.628 | 0.366 ± 0.068 | 0.26 | -0.51 |
| R1 | 71–80 | 0.658 | 0.658 | 0.670 ± 0.088 | 0.161 | 0.629 |
| R1 | 81–90 | 0.474 | 0.526 | 0.499 ± 0.255 | 0.836 | -0.169 |
| R1 | 91–100 | 0.541 | 0.541 | 0.543 ± 0.275 | 0.73 | 0.212 |
| R1 | 101–110 | 0.526 | 0.526 | 0.527 ± 0.155 | 0.836 | 0.062 |
| R1 | 111–120 | 0.464 | 0.536 | 0.383 ± 0.153 | 0.765 | -0.175 |
| R1 | 121–130 | 0.709 | 0.709 | 0.769 ± 0.201 | 0.0628 | 0.76 |
| R1 | 131–140 | 0.714 | 0.714 | 0.731 ± 0.043 | 0.0565 | 0.395 |
| R1 | 141–150 | 0.781 | 0.781 | 0.728 ± 0.112 | 0.0123 | 0.797 |
| R2 | 1–10 | 0.699 | 0.699 | 0.683 ± 0.251 | 0.0769 | 0.667 |
| R2 | 11–20 | 0.495 | 0.505 | 0.502 ± 0.352 | 0.982 | -0.016 |
| R2 | 21–30 | 0.638 | 0.638 | 0.620 ± 0.152 | 0.223 | 0.471 |
| R2 | 31–40 | 0.77 | 0.77 | 0.801 ± 0.174 | 0.0159 | 1.011 |
| R2 | 41–50 | 0.561 | 0.561 | 0.558 ± 0.246 | 0.597 | 0.17 |
| R2 | 51–60 | 0.413 | 0.587 | 0.414 ± 0.124 | 0.448 | -0.3 |
| R2 | 61–70 | 0.342 | 0.658 | 0.306 ± 0.291 | 0.161 | -0.318 |
| R2 | 71–80 | 0.342 | 0.658 | 0.381 ± 0.160 | 0.161 | -0.451 |
| R2 | 81–90 | 0.423 | 0.577 | 0.442 ± 0.258 | 0.505 | -0.36 |
| R2 | 91–100 | 0.418 | 0.582 | 0.402 ± 0.069 | 0.476 | -0.041 |
| R2 | 101–110 | 0.566 | 0.566 | 0.566 ± 0.015 | 0.566 | 0.255 |
| R2 | 111–120 | 0.577 | 0.577 | 0.607 ± 0.250 | 0.505 | 0.325 |
| R2 | 121–130 | 0.735 | 0.735 | 0.721 ± 0.278 | 0.0366 | 0.903 |
| R2 | 131–140 | 0.393 | 0.607 | 0.421 ± 0.164 | 0.346 | -0.353 |
| R2 | 141–150 | 0.311 | 0.689 | 0.297 ± 0.306 | 0.0935 | -0.693 |

**Table S8.** Comparison methods values. Values of competing methods for agreement analysis with PC2

| Sample | Batch | Label | PC2 | Normalized FrEIA | Mean DELFI | ichorCNA |
| --- | --- | --- | --- | --- | --- | --- |
| PDAC_HAD_056A | 102 | Cancer | 1.4051 | 0.9995 | 0.1707 | 0.0112 |
| CON_HAD_083A | 102 | Control | -1.4698 | -0.279 | 0.1289 | 0.0059 |
| CON_HAD_113A | 102 | Control | 1.491 | -1.0154 | 0.1558 | 0.0053 |
| PDAC_HAD_059A | 102 | Cancer | 0.2063 | 0.5914 | 0.1739 | 0.0107 |
| CON_HAD_084A | 102 | Control | -1.75 | -1.2659 | 0.1668 | 0.0074 |
| PDAC_HAD_057A | 102 | Cancer | 0.1174 | 0.9694 | 0.1782 | 0.0328 |
| PDAC_HAD_023A | 42 | Cancer | -0.1501 | 1.0325 | 0.2403 | 0.1546 |
| PDAC_HAD_012 | 42 | Cancer | 3.0679 | 0.3306 | 0.1859 | 0.0048 |
| CON_HAD_006 | 42 | Control | -0.3992 | -0.6567 | 0.1561 | 0.007 |
| CON_HAD_004 | 42 | Control | 0.5198 | -0.9853 | 0.165 | 0.0046 |
| CON_HAD_005 | 42 | Control | -0.2515 | -0.8275 | 0.1598 | 0.0045 |
| PDAC_HAD_019A | 42 | Cancer | -4.2689 | 0.8596 | 0.2404 | 0.0039 |
| PDAC_HAD_013A | 42 | Cancer | 0.4567 | 1.0218 | 0.1988 | 0.003 |
| CON_HAD_002 | 42 | Control | -0.1652 | -1.392 | 0.1442 | 0.0046 |
| PDAC_HAD_003A | 42 | Cancer | 1.9446 | 0.9005 | 0.2731 | 0.011 |
| CON_HAD_003 | 42 | Control | -3.3917 | -0.5738 | 0.1462 | 0.0076 |
| CON_HAD_001 | 42 | Control | -0.3874 | -1.0455 | 0.1461 | 0.0092 |
| PDAC_HAD_031A | 42 | Cancer | 3.025 | 1.3357 | 0.1867 | 0.0104 |
| PDAC_SHB_026 | 269 | Cancer | -0.0651 | -1.3141 | 0.1225 | 0.0058 |
| PDAC_SHB_004 | 269 | Cancer | 4.2544 | 0.9217 | 0.1913 | 0.0054 |
| CON_SHB_029 | 269 | Control | -3.8451 | -0.7468 | 0.2184 | 0.0108 |
| CON_SHB_037 | 269 | Control | -3.7985 | -0.5244 | 0.166 | 0.0093 |
| CON_SHB_036 | 269 | Control | -1.6043 | 0.1995 | 0.2048 | 0.0052 |
| CON_SHB_046 | 269 | Control | -1.9844 | -0.8449 | 0.1793 | 0.0073 |
| CON_SHB_039 | 269 | Control | -1.7959 | -0.5612 | 0.2087 | 0.0083 |
| PDAC_SHB_002 | 269 | Cancer | 3.5521 | -0.0822 | 0.1754 | 0.0099 |
| PDAC_SHB_064 | 269 | Cancer | 2.2513 | 1.1955 | 0.1986 | 0.02 |
| PDAC_SHB_024 | 269 | Cancer | 3.0356 | 1.7568 | 0.2171 | 0.003 |
| TUM_ICH_06 | 149 | Cancer | 7.7584 | 0.3046 | 0.26 | 0.0054 |
| TUM_ICH_07 | 149 | Cancer | 0.9075 | -2.2336 | 0.2097 | 0.0118 |
| CON_ICH_02 | 149 | Control | 6.7752 | 0.7187 | 0.3391 | 0.0065 |
| CON_ICH_05 | 149 | Control | -0.2871 | -0.5111 | 0.3158 | 0.0041 |
| TUM_ICH_10 | 149 | Cancer | 0.2465 | 1.3305 | 0.2466 | 0.0076 |
| TUM_ICH_09 | 149 | Cancer | -1.1818 | -0.1877 | 0.2836 | 0.0131 |
| CON_ICH_03 | 149 | Control | -1.1912 | -0.1111 | 0.2648 | 0.0071 |
| CON_ICH_06 | 149 | Control | -0.8423 | 0.0995 | 0.2463 | 0.0037 |
| CON_ICH_07 | 149 | Control | -0.1585 | 1.2883 | 0.3326 | 0.0122 |
| TUM_ICH_04 | 149 | Cancer | 2.637 | -0.1351 | 0.2639 | 0.0031 |
| TUM_ICH_03 | 149 | Cancer | -11.223 | -0.1348 | 0.2975 | 0.0052 |
| CON_ICH_01 | 149 | Control | -1.7991 | 0.5278 | 0.3169 | 0.004 |
| TUM_ICH_05 | 149 | Cancer | 2.7607 | 0.8137 | 0.2585 | 0.0042 |
| CON_ICH_09 | 149 | Control | -9.4256 | -2.0082 | 0.1931 | 0.0066 |
| CON_ICH_08 | 149 | Control | 1.836 | 0.8871 | 0.3002 | 0.0036 |
| TUM_ICH_08 | 149 | Cancer | 2.4932 | 0.2438 | 0.2238 | 0.0055 |
| TUM_ICH_01 | 149 | Cancer | 0.6944 | -0.8926 | 0.2576 | 0.0051 |

**Table S9:**
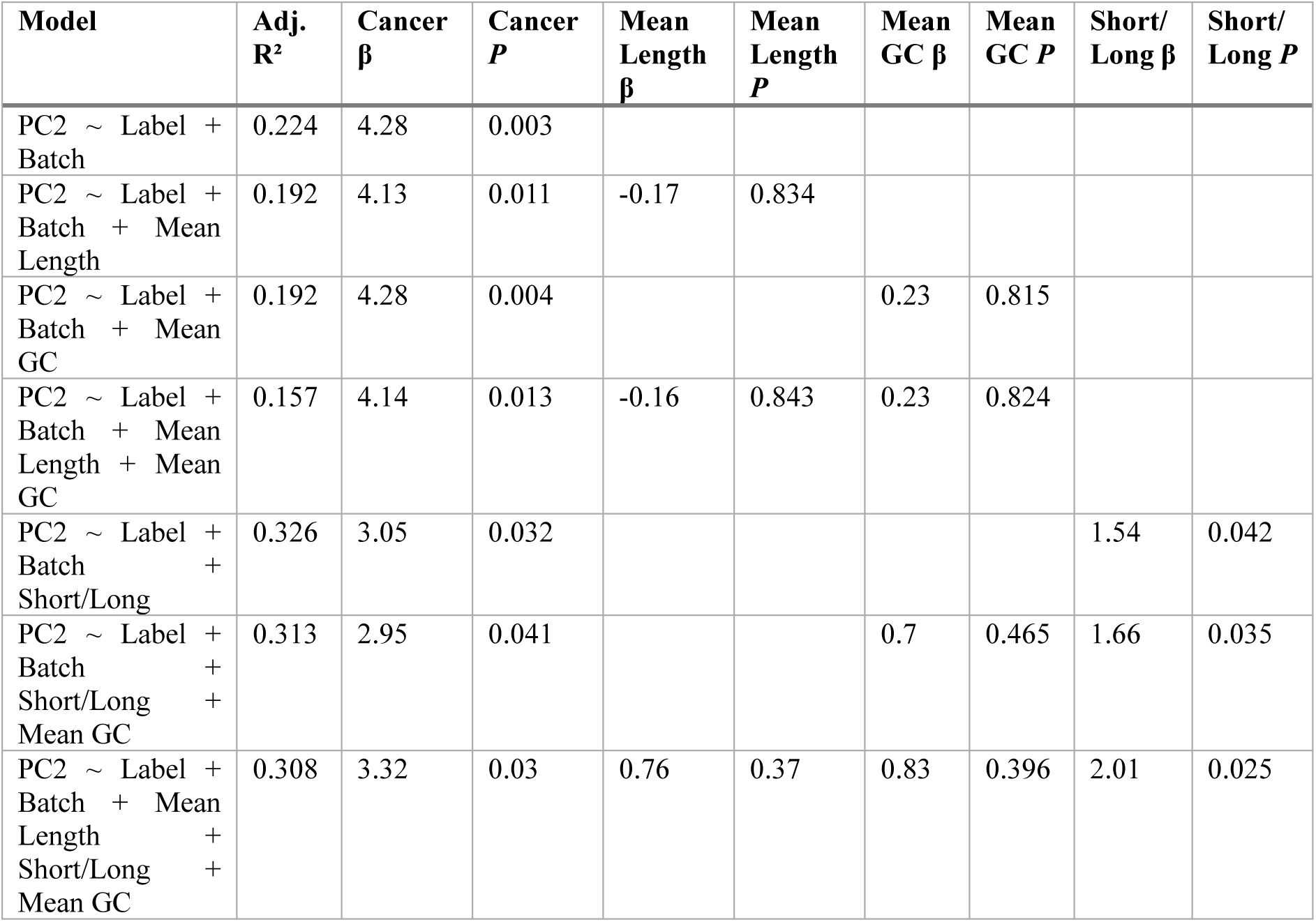
Fragment length and GC-content covariate models. Linear models evaluating whether the PBQS derived LOBO PC2 score is explained by fragment-length or GC-content covariates. The cancer/control label remained associated with PC2 after adjustment for mean fragment length and mean fragment GC content. The short-to-long fragment ratio was also associated with PC2, consistent with a fragmentomic contribution to the PBQS signal.

